# Menstruation is associated with cyclical granulysin peaks in vaginal secretions despite stable expression by cervicovaginal immune cells

**DOI:** 10.64898/2026.05.15.725524

**Authors:** Sean M. Hughes, Claire N. Levy, Dendron R. Chamberlain, Dana Varon, Britt Murphy, Katharine Schwedhelm, Juma Shafi, Jennifer M. Lund, Martin Prlic, Stephen C. De Rosa, Elizabeth Micks, Christine Johnston, Florian Hladik

## Abstract

**Problem:** The anti-microbial protein granulysin is present in vaginal secretions during the follicular phase of the menstrual cycle but nearly disappears during the luteal phase. The reason for this change is unknown.

**Method of study:** Participants (n = 23) with regular menstrual cycles collected daily vaginal swabs for granulysin ELISAs. Endocervical cytobrushes, ectocervical biopsies, vaginal biopsies, and PBMC were collected across the cycle to enumerate granulysin-expressing cells by flow cytometry. Cycle phase was determined by daily urinary luteinizing hormone testing and confirmed by serum progesterone levels.

**Results:** Granulysin levels in secretions were up to 10,000 times higher during menstruation than during the luteal phase (menstruation, median 3,924 pg/mL [IQR 400-17,280]; luteal, median and IQR undetectable [<7.81 pg/mL]). In the endocervical canal, granulysin-expressing cells were much more abundant during menstruation than during the mid-follicular or mid-luteal phases. In contrast, the number of granulysin-expressing cells in the ectocervix and vagina remained stable during the cycle. The most abundant granulysin-expressing cell types in the mucosa were CD8 T cells and NK cells. In a minority of participants, granulysin was consistently detected in luteal-phase swabs; this phenomenon was associated with parity.

**Conclusions:** Granulysin in vaginal secretions is associated with menstruation and concomitant with a spike in granulysin-expressing cells in the endocervical canal. This result explains the much higher granulysin levels in secretions during the follicular than the luteal phase. In contrast, immune cells from ectocervical and vaginal biopsies express granulysin independently of the menstrual cycle, indicating their continuous ability to respond to microbial infection.

## Introduction

Granulysin (GNLY) is an antimicrobial protein involved in cell-mediated killing of microbes^1–4^ and may play a major role in female reproductive tract immunity.^5–8^ Immune cells, especially NK and CD8 T cells, deliver GNLY and granzymes to infected cells via pore-forming perforin. Once in the cells, GNLY forms pores in microbial membranes, allowing granzymes to kill via microptosis.^4,9^ GNLY can also kill independently of perforin and granzymes: decidual NK cells transfer GNLY via nanotubes, killing intracellular microbes without killing the host cell.^5^ GNLY is synthesized as a 15 kDa protein, which is a growth factor and chemokine,^10,11^ and is cleaved to the pore-forming 9 kDa protein.^12^

The cervicovaginal tract (CVT) is the site of entry for many sexually transmitted infections (STIs) and the conduit for sperm prior to conception. It is a dynamic tissue home to many cell types, including leukocytes. Understanding CVT biology is crucial for developing mucosally-administered biomedical interventions. CVT immunity balances protection against pathogens with tolerance of foreign antigen in sperm and commensal microbiota. This balance of tolerance is adjusted in response to changing hormone levels during the menstrual cycle. Progesterone is low in the follicular phase of the menstrual cycle and rises sharply after ovulation in the luteal phase. Progesterone is thought to be immunosuppressive^13–15^ and the progesterone-high luteal phase may be a “window of vulnerability” to STIs, as the immune suppression required for embryo tolerance could reduce mucosal defenses.^16^

We recently observed a striking shift in GNLY levels in CVT secretions during the menstrual cycle: GNLY was abundant in the follicular phase and nearly absent from the luteal phase.^17^ This cyclic change in GNLY levels was among the largest observed in a meta-analysis of immune mediators in CVT secretions.^18^ However, our prior studies had limited resolution, only allowing comparison between the follicular and luteal phases, without providing finer detail within the cycle. They also lacked any information about the cellular source of the GNLY. Numerous cell types in the CVT might produce GNLY, including conventional T cells, NK cells, γδ T cells, MAIT cells, and NKT cells.^19–29^ It is unknown whether this dramatic reduction in a key antimicrobial mediator has health consequences, particularly in terms of susceptibility to STIs.

Here, we have investigated the cause of cyclical variation in cervicovaginal GNLY levels during the menstrual cycle. We measured GNLY in vaginal secretions using ELISA and enumerated GNLY-expressing cells in cervicovaginal biopsies and cytobrushes using flow cytometry. We demonstrate that the changes in GNLY levels in secretions are associated with menstruation rather than ovulation. The rise in GNLY in secretions during menstruation coincides with a dramatic increase in GNLY-expressing cells in the endocervical canal, likely derived from menstrual blood. In contrast, we demonstrate that immune cells from ectocervical and vaginal tissue biopsies express equivalent levels of GNLY during menstruation, mid-follicular phase, and luteal phase. Thus, throughout the menstrual cycle, intratissue cervicovaginal immune cells remain equally capable of expressing GNLY to respond to microbial infection.

## Methods

### Participants

Participants were eligible for this study if they were age 18 or older, premenopausal, and had regular menstrual cycles 21-35 days in length. Exclusion criteria included use of hormonal contraception, other hormonal products, or IUD within the past 2 months (6 months for injectable hormonal products); current vaginitis symptoms, gonorrhea, chlamydia, or trichomoniasis infection; active vaginal HSV-2 lesions; or positive for BV by wet mount. Wet mount was performed at the screening visit and vaginal swabs were tested for chlamydia and gonorrhea at the screening visit by Nucleic Acid Amplification Testing (Aptima). Vaginal smears were tested for bacterial vaginosis by Nugent score at all visits. If participants had evidence of a vaginitis or infection, they were treated and could be rescreened when treatment completed. All participants signed informed consent for study procedures. The study was approved by the University of Washington Human Subjects Division (STUDY00015183).

### Study design

The main study consisted of three visits (**Figure 1A**). The screening visit occurred in the mid-follicular phase, 5-9 days after the start of menstruation and at least 1 day after the end of menstruation. Starting at the screening visit, participants took daily urinary ovulation tests (Clearblue Digital Ovulation Predictor Kit) and recorded the date of the LH peak. Days prior to the LH peak were defined as the follicular phase, while days after the date of the LH peak were defined as the luteal phase. Participants attended two further visits: (1) a luteal phase visit 5-8 days after luteinizing hormone (LH) peak and (2) a follicular phase visit 5-9 days after the start of menstruation and at least 1 day after the end. The cycle phase for visits was confirmed by serum progesterone testing, with levels of <1 ng/mL for follicular and >3 ng/mL for luteal phases.

**Figure 1.**
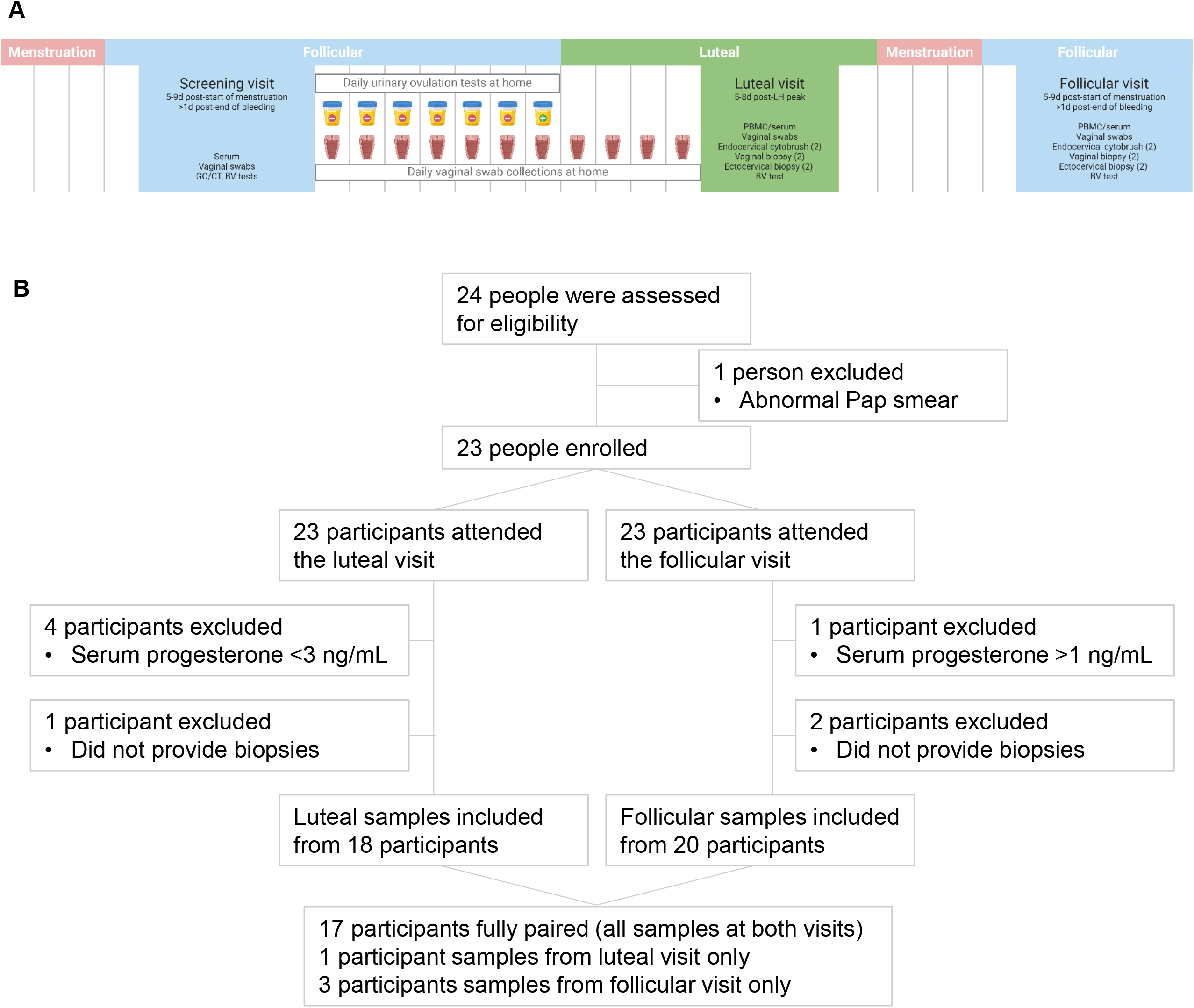
Study design and participant flow diagram. **A** Overview of clinical study design. Participant flow diagram.

### Sample collection and storage

At the luteal and follicular phase visits, participants provided 1 vaginal swab (FLOQSwabs, Copan Diagnostics), 2 endocervical cytobrushes (Digene HC2 cytobrush, Qiagen), and 4 biopsies (2 ectocervical and 2 vaginal; collected using a baby Tischler forceps), in that order (**Figure 1A**). Blood for serum and PBMC was also drawn. Cells were isolated from the endocervical cytobrushes as previously described.^22^ PBMC were isolated from blood by density gradient centrifugation. Cytobrush cells, PBMC, and biopsies were cryopreserved with 10% DMSO in fetal bovine serum and stored in a liquid nitrogen freezer.^23,30^ Swabs and serum were stored at -80°C.

### Daily at-home swabs

All participants self-collected daily vaginal swabs from the mid-follicular to the mid-luteal phase, i.e., between the screening visit and the luteal visit, and stored them in their home freezers (**Figure 1A**). Participants brought the swabs to the clinic at the luteal phase visit. Menstrual cycle phase was determined relative to the days since the LH peak for all daily at-home swabs.

### Intensive studies

Two intensive studies were also performed. In the first study, five participants collected daily vaginal swabs for a complete menstrual cycle, from the first day of menstruation through the first day of menstruation in the next cycle. In the second study, five participants provided an endocervical cytobrush, ectocervical biopsy, and vaginal biopsy on day 1 or 2 of menstruation.

### Vaginal swab processing

Swabs were processed as previously described.^31^ In brief, the swabs were incubated with phosphate-buffered saline and then eluted using a 0.45 µm SPIN-X filter (Corning Life Sciences). After elution, the samples were aliquoted to avoid future freeze/thaw cycles and frozen at -80°C. All swabs from a given participant were processed in one batch.

### Sample preparation for flow cytometry

Biopsies were digested with collagenase and DNase as previously described.^22^ In brief, biopsies were thawed, weighed, washed, and minced. They were then subjected to four 30-minute digestion periods at 37°C. After each period, the tissue pieces were washed with fresh medium over a cell strainer to separate the cells from the tissue. Tissue pieces were then combined with fresh digestion medium for another round of incubation. Cells were washed and held on ice. Cryopreserved PBMC and endocervical cytobrush cells were thawed into cell culture medium and washed in phosphate-buffered saline (PBS) prior to staining. All samples from a given participant were processed in parallel on the same day.

We performed flow cytometry in two batches for the five participants who provided biopsies during menstruation. In the first batch, as part of the main study, we ran follicular- and luteal-phase cytobrushes, ectocervical biopsies, and vaginal biopsies. In the second batch, we ran cytobrushes, ectocervical biopsies, and vaginal biopsies from the menstruation sampling study alongside remaining follicular- and luteal-phase ectocervical and vaginal biopsies from the main study. No backup endocervical cytobrushes from the main study were available for the second batch.

### Assays

#### BCA

Concentration of total protein was measured in vaginal swab eluates using the Pierce BCA Protein Assay (Thermo Fisher) relative to a standard curve of purified bovine serum albumin.

#### Granulysin ELISA

The concentration of total GNLY (9 and 15 kDa) was measured in vaginal swab eluates using the Human Granulysin DuoSet ELISA (R&D Systems) in half-area 96-well clear flat-bottom plates (Corning).

#### Flow cytometry

All samples from a given donor were processed on the same day. The antibody panel was modified from OMIP-064.^32^ Cells were first stained with LIVE/DEAD Fixable Blue Dead Cell Stain (Invitrogen) at 1:600 in PBS for 20 minutes at room temperature in the dark. Cells were washed and then stained with the surface staining antibody cocktail (Table S1) in PBS with 1% bovine serum albumin (BSA) for 20 minutes at room temperature in the dark. Cells were then fixed and permeabilized using FACS Lysing Solution and Perm 2 solution (BD Biosciences). After washing, cells were stained with the intracellular staining antibody cocktail (Table S2) in PBS with 1% BSA at 4°C overnight in the dark.^33^ The next morning, cells were washed and CountBright Absolute Counting Beads (Invitrogen) were added. The samples were acquired on a FACSymphony A5 Cell Analyzer (BD Biosciences) within 36 hours. The instrument’s configuration was described previously.^34^ The flow cytometer voltages were normalized across experiments using calibration beads (Rainbow Calibration Particles [6^th^ peak] and Supra Rainbow Midrange Fluorescent Particles, SpheroTech).

### Data analysis

#### ELISA

Log10-transformed total GNLY concentrations were compared between cycle phase using linear mixed effects models, with phase as a fixed effect and participant as a random effect to account for multiple swabs per participant.

#### Flow cytometry

Cells were gated as shown in Figure S2 using FlowJo v.10.10.0 (BD Biosciences).

#### Comparisons of cellular expression of GNLY by cycle phase

For these comparisons, we used linear mixed effects models with phase as a fixed effect and participant as a random effect. Three outcomes were used: (1) log10 percentage of the cells positive for granulysin; (2) log10 number of cells expressing granulysin (plus 1 to account for samples with 0 positive cells); and (3) log10 median fluorescence intensity (log10-transformed to normalize the distribution). Models were run separately for each cell type and form of GNLY (9 or 15-kDa). Three comparisons were calculated (Menstrual vs. follicular, menstrual vs. luteal, and follicular vs. luteal) and FDR adjustment was used to account for these repeated comparisons.

#### Presence of GNLY in luteal phase vaginal swabs

We used linear regression to test for associations between demographic variables and the detection of GNLY in vaginal swabs from the luteal phase. The model outcome was log10(n + 1), where n indicates the number of post-LH peak swabs with detectable GNLY for each participant. The numbers of swabs were incremented by 1 and log10 transformed to normalize the distribution. Pregnancy history was unavailable for two individuals. Those individuals were included in the main model as an additional explanatory category of “missing”. In a sensitivity analysis, we repeated the model, omitting those two individuals.

### Data analysis and statistics packages

All statistical analyses were performed in R^35^ version 4.4.2 using RStudio version 2025.05.1 Build 513. We used the following R packages: tidyverse,^36^ plater,^37^ and lmerTest.^38^

## Results

### Study cohort

#### Study design

We enrolled 23 participants into a three-visit study as illustrated in **Figure 1A**. Every participant attended both the luteal and follicular visits for biopsy and cytobrush collection (**Figure 1B**). Samples were excluded from 1 follicular visit and 4 luteal visits due to serum progesterone levels outside the defined ranges for those visits (<1 ng/mL for follicular; >3ng/mL for luteal). Additionally, several participants did not provide biopsies. In total, we had samples from 20 participants at the follicular visit and 18 participants at the luteal visit. We had the complete sample set from both visits from 17 participants.

All 23 participants collected daily at-home vaginal swabs between the screening visit (mid-follicular phase) and the luteal visit. Participants collected a median of 16 swabs per person (range 10-23), for a total of 354 swabs. After removing swabs with no detectable protein or insufficient volume and one swab with undefined phase, we had 348 (98.3%) usable swabs for the primary analysis (total GNLY concentration by ELISA).

Baseline participant demographics are shown in **Table 1**. Most (14/21, 61%) were nulligravid and identified as women (17/22, 77%). About one-third reported use of non-hormonal contraception, primarily condoms (hormonal contraception and IUDs were exclusion criteria). Histories of bacterial vaginosis (BV) and HSV-2 were reported in one-quarter of participants. All participants tested negative for gonorrhea and chlamydia at screening. Time-varying participant characteristics are shown by visit in **Table 2**. No yeast or sperm was detected on vaginal smears at any visit. No participants reported use of douches or vaginal medications during the study. Two participants had positive BV tests by Nugent score at both biopsy visits, and a third had a positive test at one visit.

**Table 1.** Baseline participant demographics.

| n = 23 participants | n (%) |
| --- | --- |
| Ages |  |
| <30 years | 8 (35%) |
| 30-39 years | 9 (39%) |
| ≥40 years | 6 (26%) |
| Gender identity |  |
| Woman | 17 (77%) |
| Nonbinary | 3 (14%) |
| Nonbinary, gender fluid | 1 (4.5%) |
| Questioning or unknown | 1 (4.5%) |
| Missing | 1 |
| Pregnancies (Live births) |  |
| 0 (0) | 14 (67%) |
| 1 (0) | 2 (9.5%) |
| 1 (1) | 1 (4.8%) |
| 2 (0) | 1 (4.8%) |
| 4 (2) | 1 (4.8%) |
| 5 (2) | 1 (4.8%) |
| 9 (5) | 1 (4.8%) |
| Missing | 2 |
| Use of non-hormonal contraception | 8 (35%) |
| Contraceptive method |  |
| Condom | 5 (63%) |
| Condom/withdrawal | 1 (13%) |
| Rhythm method | 1 (13%) |
| Vasectomy | 1 (13%) |
| Not applicable (no method) | 15 |
| History of HSV-2 (self-report) | 5 (22%) |
| History of BV (self-report) |  |
| Yes | 6 (26%) |
| No | 14 (61%) |
| Don't know or prefer not to answer | 3 (13%) |
| History of douche use |  |
| Yes | 2 (8.7%) |
| No | 20 (87%) |
| Don't remember | 1 (4.3%) |

**Table 2.**
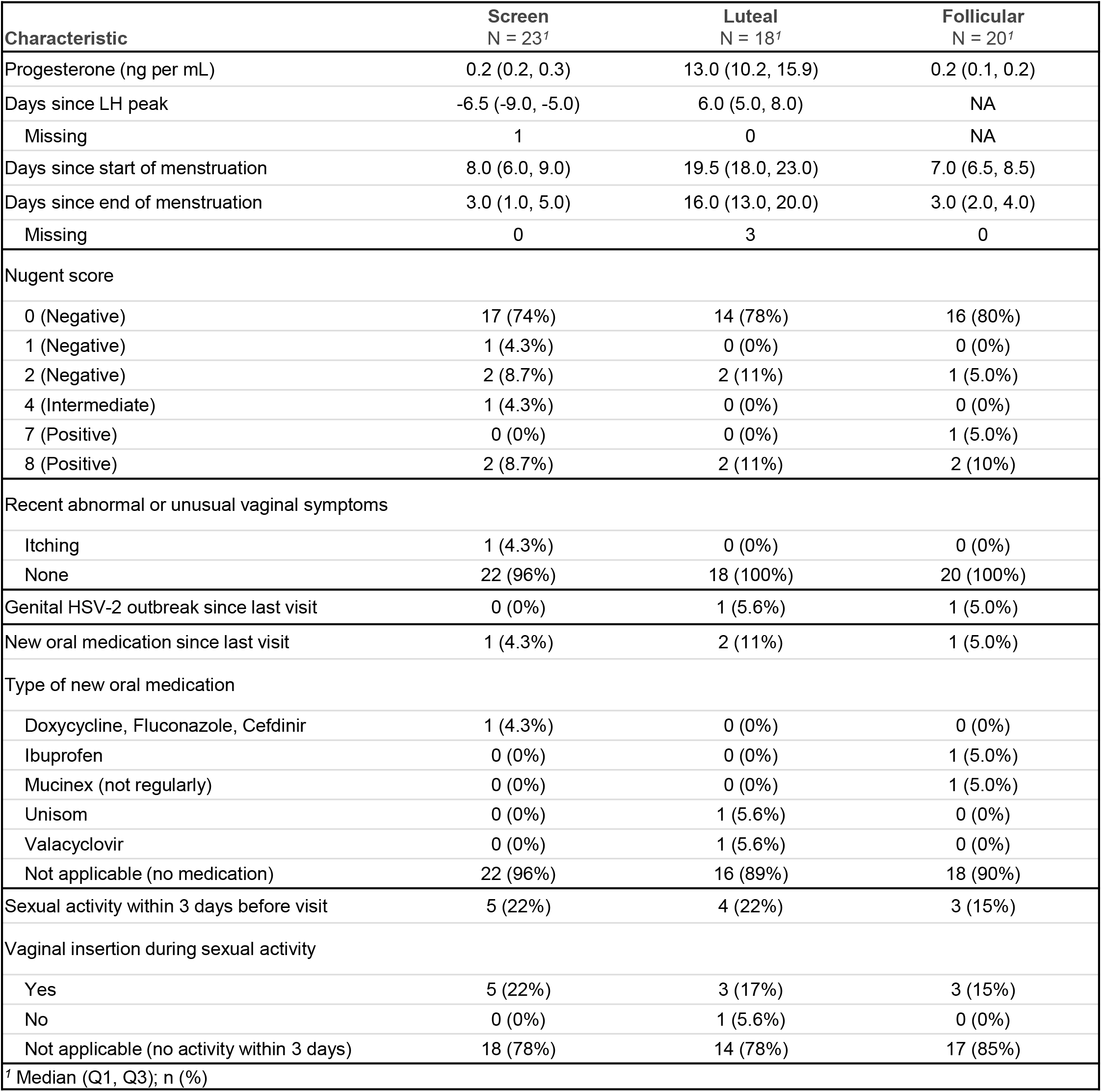
Participant characteristics by visit.

### Granulysin in vaginal secretions

#### GNLY is primarily present in vaginal secretions during the follicular phase

We first confirmed our prior findings^17,18^ of higher vaginal GNLY levels in the follicular phase than the luteal phase (**Figure 2A**). Total GNLY was detectable in more than half of follicular-phase swabs (55.7%) but less than a quarter of luteal-phase swabs (23.8%). The concentration of GNLY was substantially greater in follicular phase swabs (median 17.2 pg/mL, IQR 7.8-96.1) than in LH peak swabs (median 7.8, IQR 7.8-41.1, p = 0.002) or luteal phase swabs (median 7.8, IQR 7.8-7.8, p = 2×10^-13^).

**Figure 2.**
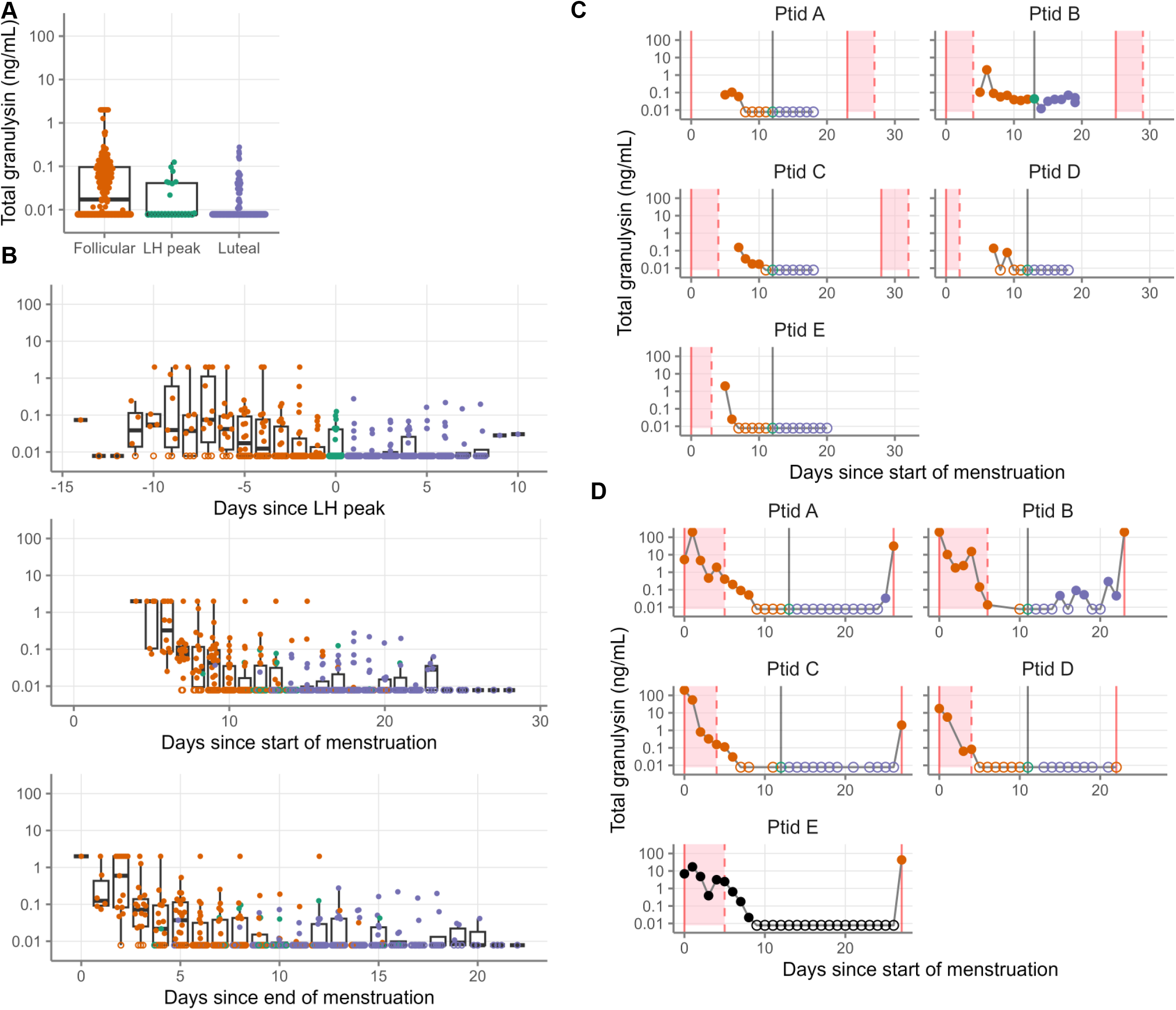
Total granulysin levels in vaginal secretions during the menstrual cycle. **A** Granulysin levels in vaginal secretions by menstrual phase, as measured by ELISA, and determined by days since LH peak (orange, follicular; green, LH peak; purple, luteal). Granulysin levels in vaginal secretions measured by ELISA, by days since start of menstruation (top), end of menstruation (middle), or LH peak (bottom). Date of end of menstruation was unavailable for 17 swabs and date of LH peak was unavailable for 21 swabs, so those swabs are not shown. **C** Daily granulysin levels in vaginal secretions measured by ELISA between the mid-follicular and mid-luteal phase for five selected participants. **D** Daily granulysin levels in vaginal secretions measured by ELISA for a separate, full menstrual cycle for the same five participants as shown in C. For all plots, filled symbols indicate detectable granulysin, and open symbols indicate below the limit of detection (0.016 ng/mL). Samples below the limit of detection were set to half the lower limit of detection (0.008 ng/mL). For A, B, and C, samples above the upper limit of detection (1 ng/mL) were set to 2 ng/mL. For D, samples above the upper limit of detection were re-run at 10- and 100-fold dilutions; samples above the upper limit of detection at 100-fold dilution were set to 200 ng/mL. For C and D, menstruation is shown in the pink-shaded regions, with the vertical pink lines showing the start (solid) and end (dashed) of menstruation. Vertical grey lines show the LH peak. Symbols are shown in black for one donor for whom the date of the LH peak was unavailable.

To confirm that this effect was due to changes in GNLY, rather than changes in total protein levels collected, we normalized GNLY concentration to total protein for each swab (**Figure S1A**). We observed a similar pattern, with the normalized concentration of GNLY substantially greater in follicular phase swabs than in LH peak (p = 0.02) or luteal phase swabs (p = 6×10^-12^).

We hypothesized that the cyclical changes in vaginal GNLY levels were due to the hormonal changes occurring around ovulation. We plotted GNLY concentrations against the number of days since LH peak, expecting to see a sharp decrease after ovulation. We did the same for days since the beginning and end of menstruation (**Figure 2B**). To our surprise, GNLY fell quickly after the end of menstruation rather than after ovulation. The same pattern was present for each individual donor (**Figure 2C** shows 5 participants, **Figure S1B** shows all participants).

#### GNLY in vaginal secretions is highest during menstruation

To confirm the association of GNLY levels with menstruation, we performed an intensive sampling study, with five previously-enrolled participants collecting a new set of daily vaginal swabs from the first day of menstruation through the first day of the next cycle. As shown in **Figure 2D**, GNLY concentrations were high throughout menstruation in all five participants, waned quickly after the end of menstruation, and then remained low until the beginning of the next cycle. Thus, we confirmed that GNLY is highest in vaginal secretions during menstruation.

### Granulysin expression by cervicovaginal cells

#### GNLY is expressed directly *ex vivo* by numerous cell types throughout the CVT

Having demonstrated the menstrual origin of GNLY in vaginal secretions, we next wanted to determine whether local production of GNLY by cervicovaginal cells was hormonally regulated. Endocervical cytobrushes, ectocervical biopsies, vaginal biopsies, and PBMC from the mid-follicular and mid-luteal phase visits were processed for flow cytometry and gated as shown in **Figure S2**. We assessed GNLY production first in CD19^-^, scatter-gated lymphocytes (**Figure S2D**, rightmost panel) and subsequently in specific cell populations (**Figures S2E and S2G**, red boxes).

GNLY was expressed *ex vivo* by lymphocytes in all specimen types in the absence of stimulation (**Figure 3A**). GNLY was detectable by two antibody clones (RB1 and DH2) which bind to different forms of GNLY and do not compete for binding. The RB1 antibody recognizes both the active 9- and the precursor 15-kDa forms of GNLY, while the DH2 antibody recognizes only the 9-kDa form.^5^ Thus, we designate RB1^+^DH2^-^ single-positive cells as expressing “GNLY15” and RB1^+^DH2^+^ double-positive cells as expressing “GNLY9” (though they may also express GNLY15). The 15-kDa form alone was expressed by 2-3% of CVT lymphocytes and the 9-kDa form by about 1% (**Figure 3B**). The proportions were similar across CVT sample types, but much higher in PBMC (8-10%). Among GNLY-expressing cells, the median fluorescence intensity of GNLY was similar across sample types (**Figure 3C**). Classical CD8 T cells (pink boxes) made up the largest proportion of the GNLY15-expressing lymphocytes in the CVT, while NK cells (blue boxes) were the main expressers of GNLY9 (**Figure 3D**). In PBMC, in contrast, NK cells were the main source of both forms of GNLY.

**Figure 3.**
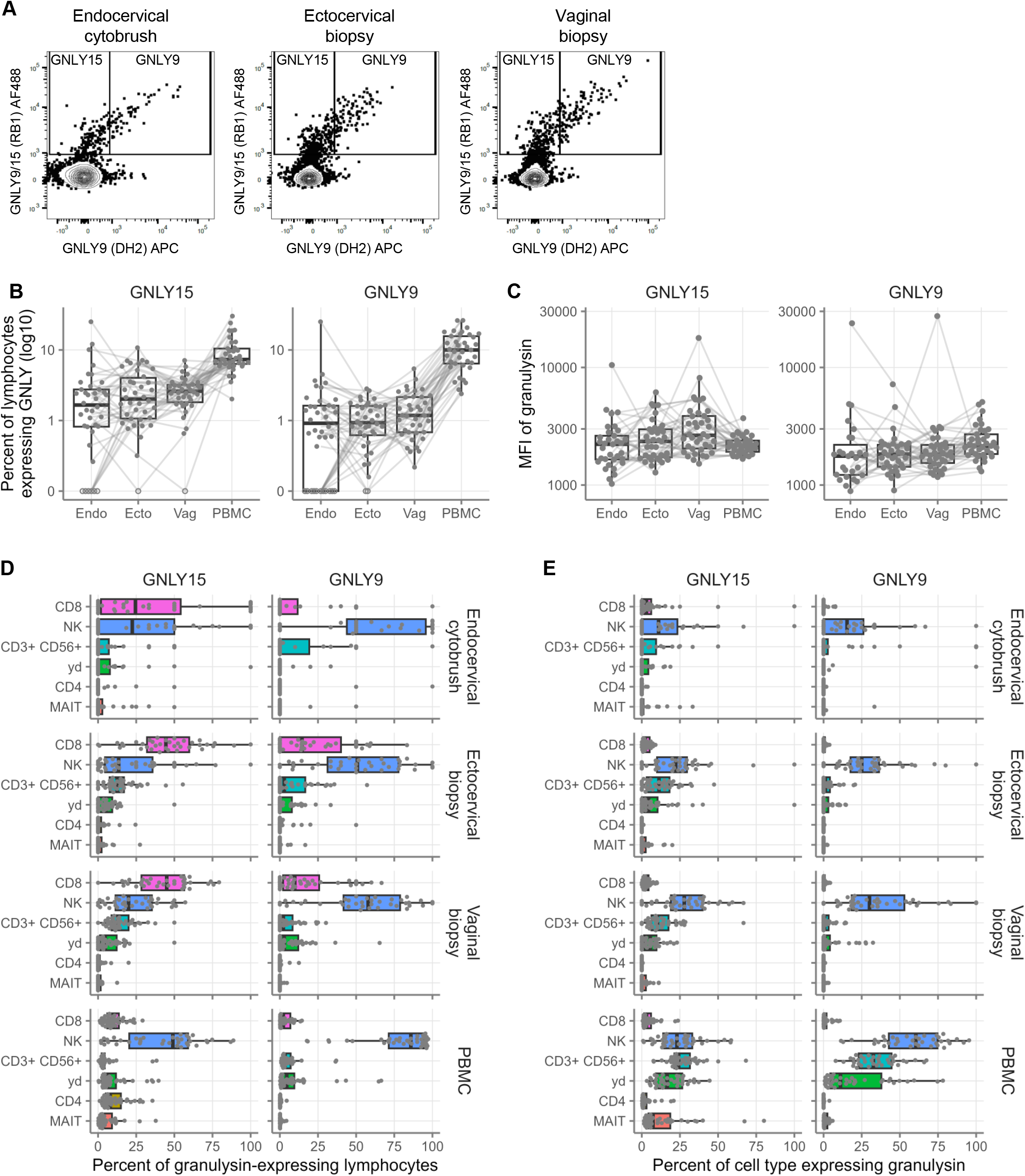
Granulysin expression by cervicovaginal lymphocytes. **A** Granulysin expression in CD19-, scatter-gated lymphocytes. Total granulysin (9 and 15-kDa forms) is shown on the y-axis, and the 9-kDa form on the x-axis. The samples shown are concatenations of all samples from all donors of the indicated types. Percent of CD19-, scatter-gated lymphocytes expressing GNLY15 (left) and GNLY9 (right). Open symbols indicate below the limit of detection (no positive cells in that sample). The y-axis scale is modified to show samples below the limit of detection at 0 on the log scale. **C** Median fluorescence index of the RB1 antibody clone (total GNLY, left) and DH2 antibody clone (9 kDa GNLY, right) in CD19-, scatter-gated lymphocytes positive for those antibodies. **D** Phenotypes of GNLY-expressing cells. Percent of GNLY15-expressing (left) and GNLY-9-expressing (right) lymphocytes made up by the indicated cell types. **E** Percentages of each indicated cell type expressing GNLY15 (left) or GNLY9 (right).

On a per cell basis, GNLY, and especially GNLY9, was expressed at the highest rates in NK cells (**Figure 3E**). However, NK cells were quite rare in the mucosal samples compared to CD8 T cells. Because of that, most GNLY15-expressing cells in the CVT were CD8 T cells.

### Menstrual cycle-related variation in granulysin expression by cervicovaginal cells

#### Cellular expression of GNLY was much higher during menstruation in cytobrushes but steady in all other sample types, regardless of cycle phase

Given the menstrual origin of GNLY in vaginal secretions, we conducted an intensive study collecting samples on day 1 or 2 of menstruation in 5 participants, four of whom were previously enrolled in the main study. The combined results from the main study (mid-follicular and mid-luteal) and this follow-up study (menstruation) are shown in **Figure 4** with statistical results shown in **Tables S3-5**. Data from all visits with concordant serum progesterone levels from the main study are included in this analysis (n=20 follicular, n=18 luteal, with n=17 paired) as well as from the 5 participants who provided samples during menstruation. The samples from the follow-up menstruation study were collected and processed separately from the main study.

**Figure 4.**
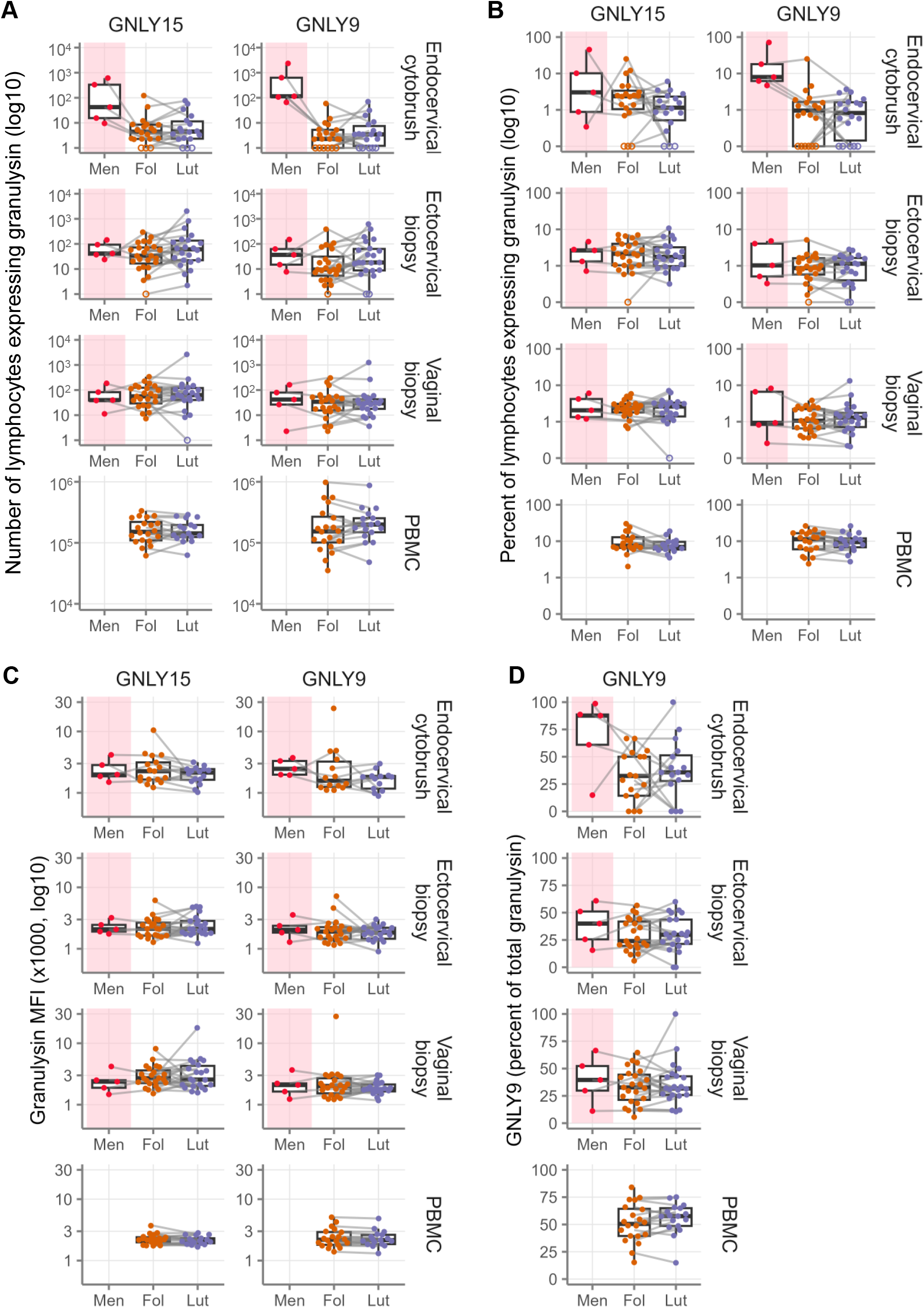
Granulysin expression by cervicovaginal lymphocytes stratified by menstrual cycle phase. **A** Number of CD19-, scatter-gated lymphocytes expressing GNLY15 (left) and GNLY9 (right) in the indicated specimens in the menstrual, follicular, and luteal phases. The y-axis scale is modified to show samples below the limit of detection at 0 on the log scale. Percent of CD19-, scatter-gated lymphocytes expressing GNLY15 (left) and GNLY9 (right) in the indicated specimens in the menstrual, follicular, and luteal phases, as in A. **C** Median fluorescence index of the RB1 antibody clone (total GNLY, left) and DH2 antibody clone (9 kDa GNLY, right) in the indicated specimens in the menstrual, follicular, and luteal phases. **D** Percent of GNLY-expressing CD19-, scatter-gated lymphocytes that express GNLY9. Menstrual is shown in red, follicular is shown in orange, and luteal is shown in purple. Open symbols indicate below the limit of detection (no positive cells in that sample). Paired samples from the same donor are connected by lines.

Dramatically more lymphocytes from endocervical cytobrushes expressed GNLY during menstruation than in the mid-follicular or mid-luteal phase visits (**Figure 4A**, **Table S3**). In addition, a much larger percentage expressed GNLY9 (**Figure 4B**, **Table S4**). In contrast, GNLY^+^ cells were not more abundant during menstruation for the other sample types (ectocervical biopsies, vaginal biopsies, or PBMC). In addition, GNLY production was similar between the follicular and luteal phases for all sample types. The MFI of both total GNLY (antibody clone RB1) in RB1-positive cells as well as the MFI of GNLY-9 (antibody clone DH2) in DH2-positive cells were consistent across the cycle (**Figure 4C**, **Table S5**). Finally, we observed a shift in granulysin expression from GNLY 15 to GNLY9 in cytobrushes during menstruation (**Figure 4D**). Thus, cellular expression of GNLY was much higher during menstruation in cytobrushes but steady in all other sample types, regardless of cycle phase. This result is consistent with menstrual fluid being the source of most cervicovaginal GNLY.

We next sought to determine the cell type responsible for the elevated levels of GNLY expression in endocervical cytobrushes during menstruation. To do so, we analyzed the six GNLY-expressing cell types from Figure 3D and E in cytobrushes. The number of GNLY-expressing cells increased substantially during menstruation for all six cell types (**Figure 5A**, **Table S6**). NK cells were the most abundant GNLY-expressing cell type during menstruation for both forms of GNLY. The percentage of cells expressing GNLY also increased during menstruation for most of these cell types, in particular for GNLY9 (**Figure 5B**, **Table S7**). This result suggests that multiple cell types contribute to the rise in GNLY production during menstruation, with NK cells contributing the most.

**Figure 5.**
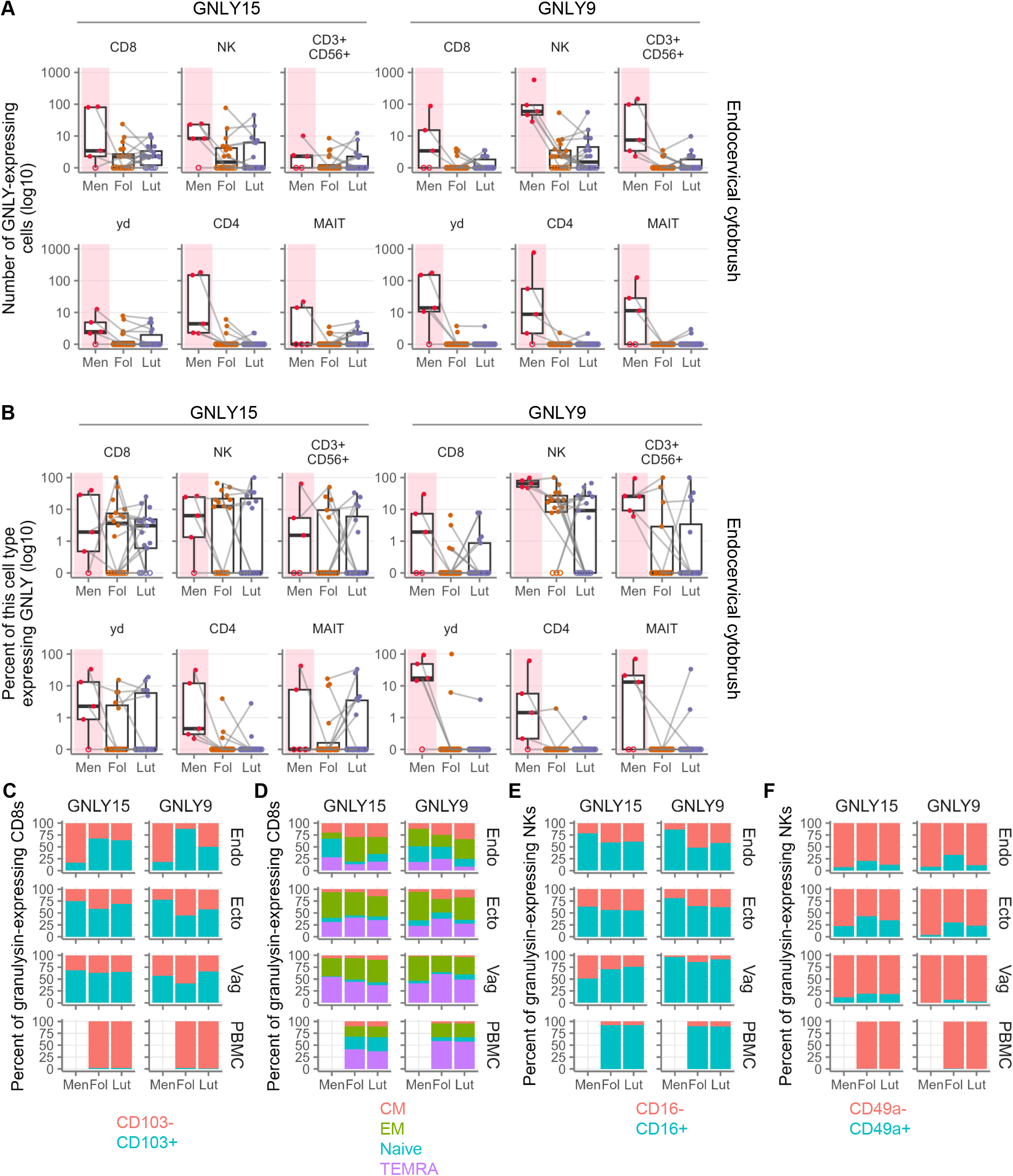
Granulysin expression by subtypes of cervicovaginal cells stratified by menstrual cycle phase. **A** Number of the indicated cell types expressing GNLY15 (left) and GNLY9 (right) in endocervical cytobrushes in the menstrual, follicular, and luteal phases. The y-axis scale is modified to show samples below the limit of detection at 0 on the log scale. Menstrual is shown in red, follicular is shown in orange, and luteal is shown in purple. Open symbols indicate below the limit of detection (no positive cells in that sample). Paired samples from the same donor are connected by lines. Percent of the indicated cell types expressing GNLY15 (left) and GNLY9 (right) in the endocervical cytobrushes in the menstrual, follicular, and luteal phases, as in A. **C** Mean percentage of CD8 T cells expressing CD103. **D** Mean percentage of CD8 T cells with each memory phenotype. **E** Mean percentage of NK cells expressing CD16.

#### GNLY-expressing cells from the endocervical canal resemble peripheral cells

We further phenotyped the GNLY-expressing NK cells and CD8 T cells from the endocervical cytobrushes to determine their likely origin. To our surprise, we found that their phenotypes were more suggestive of an origin in peripheral blood than an endometrial or cervicovaginal-resident origin. Specifically, like those in PBMC, the GNLY-expressing CD8 T cells in the endocervix during menstruation mostly lacked the tissue residency marker CD103, in contrast to other times during the cycle (**Figure 5C**). GNLY-expressing CD8 T cells from vaginal and ectocervical biopsies had consistent levels of CD103 expression throughout the cycle. Similarly, a substantial proportion of the endocervical menstrual phase GNLY-expressing CD8 T cells had a naïve phenotype, like in the blood, and unlike elsewhere in the CV tract and other times during the cycle (**Figure 5D**). Endocervical menstrual phase GNLY-expressing NK cells expressed CD16 at especially high rates, again like the blood (**Figure 5E**). The uterine NK cell marker CD49a was rare on all cervicovaginal NK cells, but especially GNLY-expressing endocervical NK cells during menstruation (**Figure 5F**). These results, and those shown in Figure 4D, are consistent with increased GNLY in the endocervical canal during menstruation coming from peripheral-like immune cells, rather than from local or endometrial-like immune cells.

### Granulysin in luteal-phase vaginal secretions

#### Presence of granulysin in luteal-phase swabs is associated with parity

GNLY was undetectable in most, but not all, luteal phase swabs, raising the question of what causes GNLY production during the luteal phase. We stratified participants into 3 groups: those with no GNLY in luteal phase swabs (n=9), those with GNLY in 1-2 luteal swabs (n = 8), and those with GNLY in 5-7 luteal swabs (n=4, **Figure 6A**). In one individual for whom data were available for two cycles, we observed GNLY in numerous luteal-phase swabs in both cycles (**Figure 6B**), suggesting this is a consistent phenomenon.

**Figure 6.**
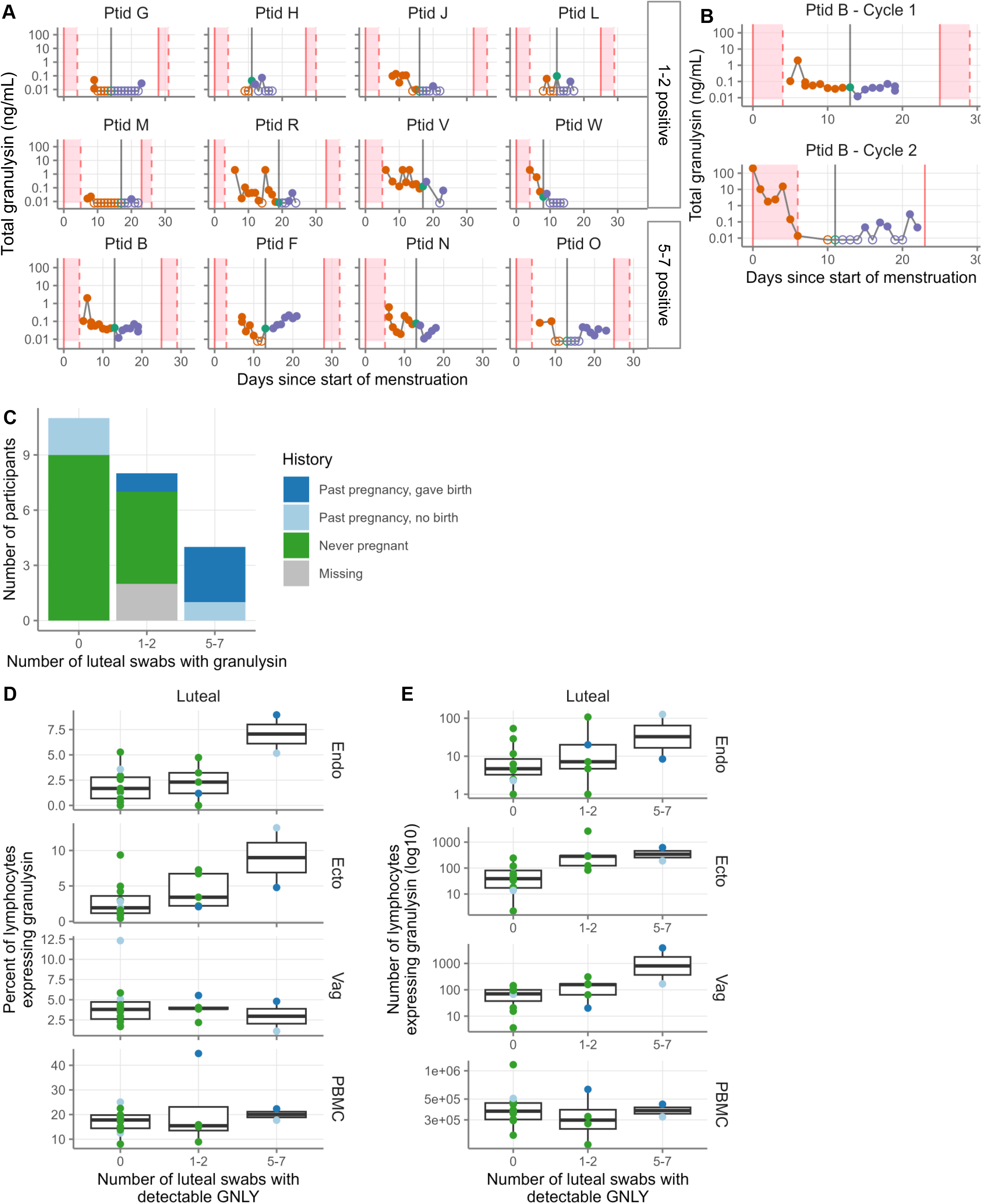
Expression of granulysin in luteal phase swabs. **A** Daily granulysin levels in vaginal secretions measured by ELISA between the mid-follicular and mid-luteal phase for participants who had 1-2 positive swabs in the luteal phase (top two rows) or 5-7 positive swabs in the luteal phase (bottom row). Daily granulysin levels in vaginal secretions measured by ELISA for one participant over two different cycles. **C** Gravidity and parity stratified by number of luteal swabs positive for granulysin. **D** Percent of CD19-, scatter-gated lymphocytes expressing GNLY in the luteal phase in participants with 0, 1-2, or 5-7 positive swabs in the luteal phase. Symbols are colored by gravidity and parity as in C. **E** Number of CD19-, scatter-gated lymphocytes expressing GNLY in the luteal phase in participants with 0, 1-2, or 5-7 positive swabs in the luteal phase. Symbols are colored by gravidity and parity as in C. Endo, endocervical cytobrush; ecto, ectocervical biopsy; vag, vaginal biopsy.

We considered several demographic variables as possible explanatory factors: participant age, HSV-2 history, BV history, use of contraception, use of lubricant during sexual activity, and gravidity/parity. In univariate linear models, parity was significantly positively associated with detection of GNLY in luteal phase swabs (p = 0.0009 in **Table S8**; and **Figure 6C**). Given the association of age with parity, we combined age and birth history into a bivariate model, where we determined that a history of giving birth was positively associated with the number of GNLY^+^ swabs in the luteal phase (p = 0.006), while age was not (p = 0.67; **Table S9**). Thus, parity is likely associated with the presence of GNLY in luteal phase swabs, but this result needs to be confirmed in a larger study designed and powered for this purpose.

Our next question was whether the luteal phase GNLY was produced locally in the CVT or elsewhere. We checked for an association between detectable luteal GNLY in vaginal swabs and the percentage and number of local GNLY-expressing lymphocytes in luteal phase samples. Cellular data were only available from two individuals with 5-7 swabs positive for GNLY, precluding statistical testing. We did observe in these individuals an increase in the percentage of GNLY^+^ lymphocytes in endocervical cytobrushes and cervical biopsies (**Figure 6D**), as well as an increased absolute number of GNLY^+^ lymphocytes in all three CVT sample types (**Figure 6E**). In contrast, levels were steady in PBMC. Thus, there may be increased local production of GNLY in the luteal phase in these individuals.

## Discussion

Here, we show a dramatic increase in GNLY in vaginal secretions during the follicular phase of the menstrual cycle, in particular during menstruation. In keeping with that result, GNLY-expressing cells were much more abundant in endocervical cytobrush samples during menstruation than during the mid-follicular or mid-luteal phase. In contrast, local immune cells from cervical and vaginal biopsies maintained consistent GNLY frequencies throughout the cycle. We further show that luteal phase secretions from some participants contained GNLY; this phenomenon was associated with parity.

Immune cells throughout the cervix and vagina expressed GNLY, particularly CD8^+^ T cells and NK cells. While NK cells expressed GNLY at higher rates, CD8^+^ T cells were more abundant and made up the largest proportion of GNLY-expressing cells in the CVT. GNLY expression was more restricted in the tissue than in blood, with much higher percentages of PBMC lymphocytes expressing GNLY than their mucosal counterparts, as has previously been shown in gut, lung, liver, and skin.^39^ However, the MFIs of positive cells were similar between tissue and blood. This suggests that tissue immune cells are less likely to express GNLY, but that individual GNLY^+^ cells express similar quantities of GNLY between the tissue and blood. The mechanisms regulating GNLY expression in the tissue are unknown. One factor may be cell type, as high percentages of CD16^+^ NK cells expressed GNLY, but this cell type was much rarer in the tissue than in the blood, as has been seen previously in the tonsil, gut, and lymph node.^40^ Similarly, expression of tissue residency markers was inversely associated with granulysin expression in CD8 T cells.^39^ Thus, a temporary influx of peripheral-like cells could dramatically alter local GNLY expression.

We expected that the large increase in GNLY expression in the endocervix during menstruation would be driven by shed endometrial cells. Surprisingly, we found that the cells were more phenotypically similar to peripheral blood cells than to endometrial immune cells. Specifically, the GNLY-expressing CD8 T cells had lower expression of the tissue residency marker CD103 and more often had a naïve phenotype. The GNLY-expressing NK cells had low expression of CD49a and high expression of CD16, unlike uterine NK cells, which express CD49a and have low CD16 expression.^41,42^ Similar to uterine NK cells, menstrual blood collected in menstrual cups contains high levels of CD56^high^CD16^-^ NK cells.^43^ Notably, we did not collect menstrual blood directly; the cytobrushes we collected during menstruation likely include a combination of cells derived from menstrual blood and local endocervical cells. The CD56^high^CD16^-^ NK cells present in menstrual blood may play a role in GNLY levels in vaginal secretions during menstruation. One possible explanation for our surprising finding of primarily peripheral-like CD49a^-^CD16^+^ NK cells in endocervical cytobrushes during menstruation is that NK cells may exhibit differential adherence to the endocervix during menstruation. Peripheral-like NK cells may be more likely to collect on the surface of the endocervical canal (and thus be obtained in a cytobrush) and uterine NK cells may be more likely to pass through (and thus be obtained in a menstrual cup).

Our data suggest that the increase in vaginal GNLY during the follicular phase derives from an influx of GNLY-expressing cells and extracellular GNLY during menstruation. In our previous meta-analysis of cervicovaginal immune modulators, we found that among those factors that changed across the cycle, most were lower in the luteal phase, similar to what we observed for GNLY.^18^ Thus, the present study raises the possibility that the cervicovaginal changes in some of those factors are a consequence of menstruation, rather than changes to production in the cervix or vagina. For instance, matrix metalloproteinase (MMP)-1 and MMP-3 both decreased in the luteal phase.^18^ MMPs are key proteinases involved in the breakdown of the endometrial extracellular matrix during menstruation.^44^ The large decrease in MMP-1 and MMP-3 during the luteal phase may, like GNLY, reflect menstrual shedding of these factors rather than changes in cervicovaginal production. Furthermore, we find that traces of menstruation can be detected well beyond the reported end of menstrual bleeding, with granulysin detected several days past that point in nearly all participants, albeit at a much lower concentration (**Figure S1**). Thus, changes in some immune modulators across the menstrual cycle may reflect their presence in menstrual fluid, rather than local production. However, this would need to be determined independently for each immune modulator and take into account possible differential local production throughout the cycle as well as kinetics in response to hormonal changes. Moreover, whether their origin is local or in menstrual fluid, the immune modulators are nonetheless present in vaginal secretions and may exert essential functions there.

The function of GNLY in vaginal secretions is unknown. One possible function is antimicrobial killing. In one *in vitro* experimental system, the IC50 for purified 9-kDa GNLY is 10 μg/mL,^3^ about 100-times higher than our maximum measured concentrations, suggesting that the concentration in vaginal secretions is too low to kill microbes. However, it should be noted that our concentrations are diluted by an unknown factor, given that we eluted the (variable) volume of secretions on each swab into 800 μL of saline. Thus, the true concentration of GNLY may approach the *in vitro* IC50. In addition, the potency of GNLY is much higher in the presence of granzymes.^3^ Further work is needed to determine the function of GNLY in vaginal secretions.

While GNLY was rare in vaginal secretions during the luteal phase, it was present in 5-7 swabs each from several participants. The strongest association was with parity. There are several possible explanations for this association. First, following pregnancy, the cervical os changes from pinpoint to a slit-like and more open configuration.^45^ This anatomical change might allow increased transit of GNLY from the endometrium into the vaginal canal. Second, several studies have shown a positive association between pregnancy and GNLY production. One study showed a positive correlation between gravidity and levels of GNLY in the decidua.^46^ Two other studies showed that pregnancy-trained decidual NK cells appear in the decidua of subsequent pregnancies and express higher levels of GNLY than during first pregnancies.^47,48^ It is unclear, however, whether these findings from the decidua can be extended to the non-pregnant endometrium.

Our study had several limitations. We only measured *ex vivo* GNLY expression, rather than production in response to stimulation. It is therefore possible that we missed a differential cellular capacity to respond to stimulation at different phases. While the mid-follicular and mid-luteal visits were conducted in sequential cycles in most participants, the menstrual phase samples were collected and processed months later. This batch effect could potentially confound our results. However, differences in GNLY expression during the menstrual phase were observed only in cytobrushes; the ectocervical and vaginal biopsies from the menstrual phase closely matched the results of the mid-follicular and mid-luteal phase biopsies. In addition, the mid-follicular and mid-luteal phase biopsies were run in both batches and gave similar results regardless of batch. These controls suggest that the difference we observed in cytobrushes was real. Several possible confounding factors may have influenced the measured granulysin concentrations or cellular phenotypes, including cyclical differences in cervical mucus and the efficiency of collection of vaginal secretions as well as sample cryopreservation, sexual activity, and microbiome composition. The two intensive studies both included only 5 participants, so the conclusions about menstruation in particular are drawn from a small sample size. Our description of an influx of “peripheral-like” cells during menstruation is based on phenotypic markers, rather than a direct demonstration of actual transit from the periphery. Only four participants had more than two swabs with GNLY present in the luteal phase, which weakens our ability to discern potential causes of luteal phase GNLY. Finally, cell numbers were low in our mucosal samples, which prevented us from conducting very fine phenotyping.

In summary, we demonstrate a striking increase in GNLY concentrations in vaginal secretions during the follicular phase of the menstrual cycle, especially during menstruation. We show that an influx of peripheral-like GNLY-expressing lymphocytes, especially NK cells, is associated with a large increase in GNLY production in the endocervix during menstruation. In contrast, GNLY expression by local immune cells is largely steady between the follicular and luteal phases of the cycle. We additionally show that the presence of GNLY in secretions during the luteal phase was associated with parity. Thus, while GNLY levels in secretions change dramatically during the menstrual cycle, expression by local mucosal cells remains steady throughout the cycle.

## Acknowledgments

We would like to thank the study participants for their time and participation; the staff of the VRC, especially Kirsten Hauge, Matt Seymour, Gina Huynh, and Chloe Wilkens, for performing the clinical studies; and Ronit Katz for statistical expertise and consulting. FLOQSwabs were provided free of charge by the manufacturer (Copan Diagnostics).

## Ethics statement

All participants signed informed consent for study procedures. The study was approved by the University of Washington Human Subjects Division (STUDY00015183).

---

**Figure S1.**
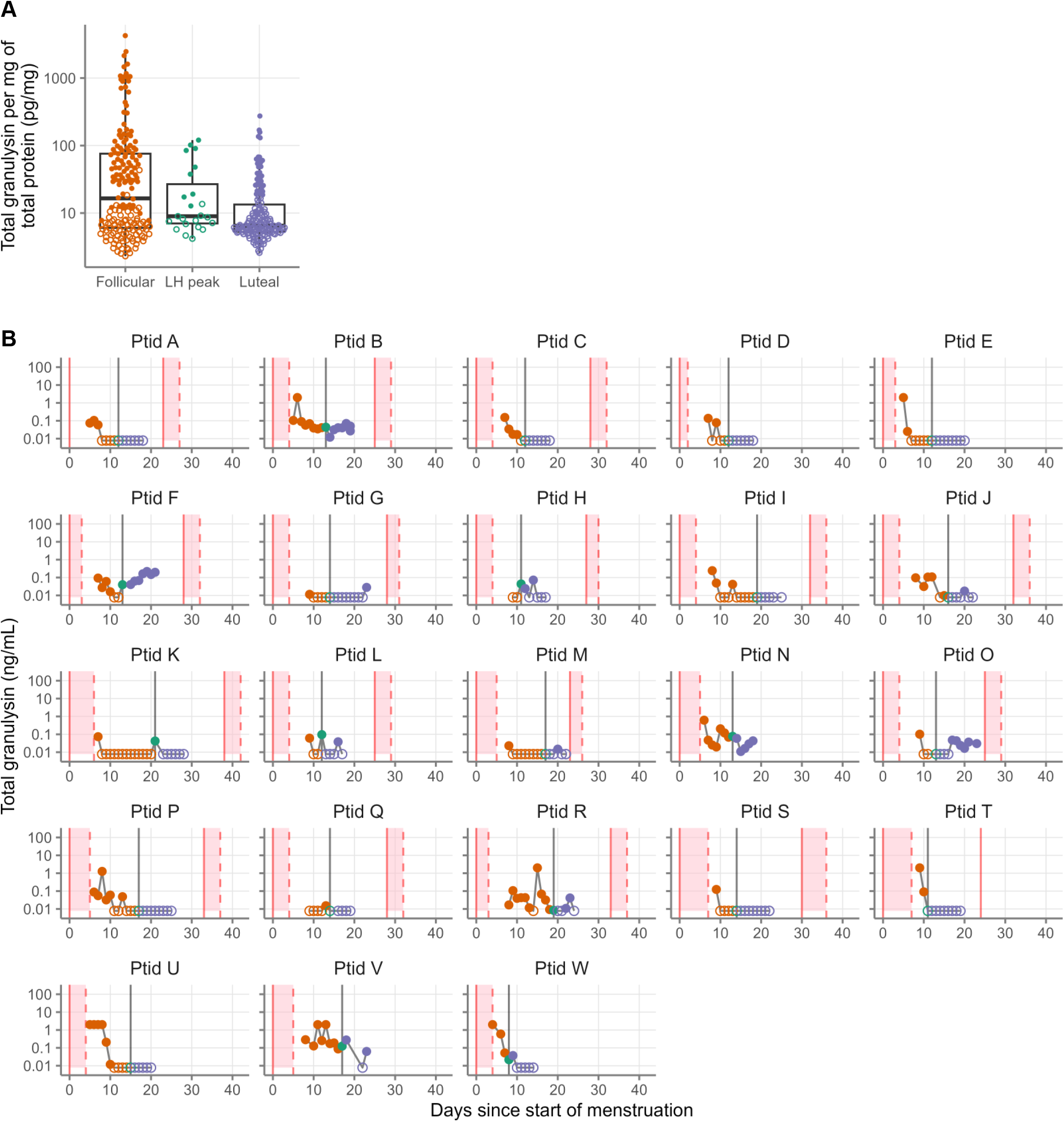
Total granulysin levels in vaginal secretions during the menstrual cycle. **A** Granulysin levels in vaginal secretions measured by ELISA normalized to total protein levels measured by BCA assay compared to menstrual phase as determined by days since LH peak. Daily granulysin levels in vaginal secretions measured by ELISA between the mid-follicular and mid-luteal phase for all participants. For all plots, filled symbols indicate detectable granulysin and open symbols indicate below the limit of detection (0.016 ng/mL). Samples below the limit of detection were set to half the lower limit of detection (0.008 ng/mL). Samples above the upper limit of detection (1 ng/mL) were set to 2 ng/mL. For B, menstruation is shown in the pink-shaded regions, with vertical pink lines indicating the start (solid) and end (dashed) of menstruation. Vertical grey lines show the LH peak. Symbols are shown in black for one donor for whom the date of the LH peak was unavailable.

**Figure S2.**
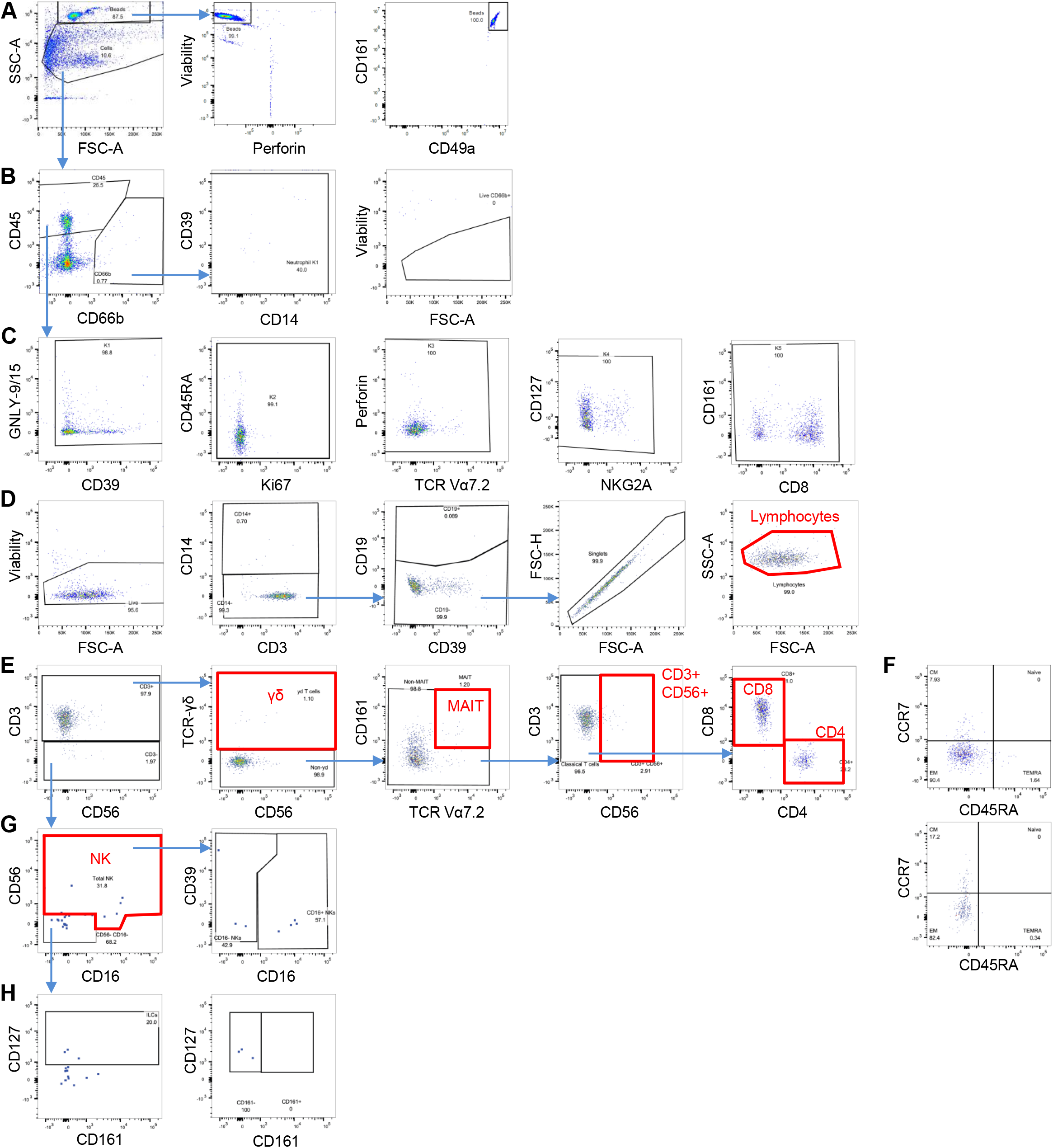
Gating scheme using a representative vaginal cell sample. Populations used in later analyses are highlighted in red. **A** Gating of counting beads (continues at right) and cells (continues in B). Identification of neutrophils (continues at right with removal of autofluorescence, debris and fluorescent aggregates and viability gating) and CD45+ non-neutrophils (continues in C). **C** Removal of autofluorescence, debris and fluorescent aggregates by exclusion of events outside the normal range for the indicated markers to obtain a clean cell population (continues in D). **D** Identification of live, CD14-, CD19-, singlet lymphocytes (continues in E). **E** Phenotyping of γδ T cells, mucosal-associated invariant T (MAIT) cells, CD3+ CD56+ cells, and classical CD4 and CD8 T cells. **F** Continuing from E, memory phenotyping of classical CD8 (top) and CD4 (bottom) T cells. **G** Continuing from E, identification and phenotyping of NK cells. The symbols are enlarged for clarity because they are rare. **H** Continuing from G, identification and phenotyping of innate lymphoid cells (ILCs). The symbols are enlarged for clarity because they are rare.

**Table S1.** Extracellular flow cytometry antibody panel.

| Laser | Detector | Filter | Antigen | Fluorochrome | Company | Catalog number | Clone | RRID | Titer per 100 uL |
| --- | --- | --- | --- | --- | --- | --- | --- | --- | --- |
| UV (355 nm) | U395 | 379/28 | CD45 | BUV395 | BD | 563792 | HI30 | AB_2869519 | 0.1 |
| UV (355 nm) | U500 | 515/30 | CD3 | BUV496 | BD | 612940 | UCHT1 | AB_2870222 | 0.5 |
| UV (355 nm) | U570 | 570/40 | CD19 | BUV563 | BD | 612916 | SJ25C1 | AB_2870201 | 0.1 |
| UV (355 nm) | U730 | 740/35 | CD103 | BUV737 | BD | 563850 | Ber-ACT8 | AB_3676338 | 0.2 |
| UV (355 nm) | U780 | 785/62 | CD8 | BUV805 | BD | 612889 | SK1 | AB_2833078 | 0.1 |
| Violet (405 nm) | V510 | 515/20 | CD4 | BV480 | BD | 566104 | SK3 | AB_2739506 | 0.05 |
| Violet (405 nm) | V570 | 575/25 | CD16 | BV570 | Biolegend | 302036 | 3G8 | AB_2632790 | 0.3 |
| Violet (405 nm) | V610 | 605/20 | CCR7 | BV605 | BD | 566754 | 2-L1-A | AB_2869850 | 10 |
| Violet (405 nm) | V650 | 660/20 | NKG2D | BV650 | BD | 563408 | 1D11 | AB_2738188 | 2.5 |
| Violet (405 nm) | V750 | 750/30 | CD56 | BV750 | Biolegend | 362556 | 5.1H11 | AB_2801001 | 0.5 |
| Violet (405 nm) | V780 | 785/62 | TCR Va7.2 | BV785 | Biolegend | 351722 | 3C10 | AB_2566042 | 2.5 |
| Blue (488 nm) | B710 | 710/50 | CD66b | RB705 | BD | 570599 | G10F5 | AB_3685879 | 0.5 |
| Blue (488 nm) | B780 | 780/40 | CD127 | RB780 | BD | 568750 | HIL-7R-M21 | AB_3684515 | 5 |
| Green (561 nm) | G575 | 575/25 | CD39 | PE | BD | 567156 | A1 | AB_2916473 | 0.5 |
| Green (561 nm) | G610 | 610/20 | TCR-$\gamma\delta$ | PE-CF594 | BD | custom | 11F2 | AB_400356 (purified) | 1.5 |
| Green (561 nm) | G710 | 710/50 | CD14 | PE-Fire700 | Biolegend | 367158 | 63D3 | AB_2876695 | 0.25 |
| Green (561 nm) | G780 | 780/60 | NKG2A | PE-Cy7 | Coulter | B10246 | Z199 | AB_2687887 | 0.1 |
| Red (640 nm) | R710 | 730/45 | CD45RA | R718 | BD | 567073 | HI100 | AB_2916420 | 0.25 |
| Red (640 nm) | R780 | 780/60 | CD161 | APC-Fire750 | Biolegend | 339944 | HP-3G10 | AB_2617016 | 5 |

**Table S2.** Intracellular flow cytometry antibody panel.

| Laser | Detector | Filter | Antigen | Fluorochrome | Company | Catalog number | Clone | RRID | Titer per 100 uL |
| --- | --- | --- | --- | --- | --- | --- | --- | --- | --- |
| Violet (405 nm) | V450 | 450/40 | Perforin | BV421 | BD | 563393 | δG9 | AB_2738178 | 0.3 |
| Blue (488 nm) | B515 | 515/20 | GNLY-9/15 RB1 | AF488 | BD | 558254 | RB1 | AB_2869123 | 0.625 |
| Blue (488 nm) | B660 | 660/40 | Ki67 | BB660 | BD | custom | B56 | AB_396287 (purified) | 0.02 |
| Red (640 nm) | R660 | 670/30 | GNLY-9 DH2 | APC | Biolegend | 348010 | DH2 | AB_2819989 | 1.25 |

**Table S3.**
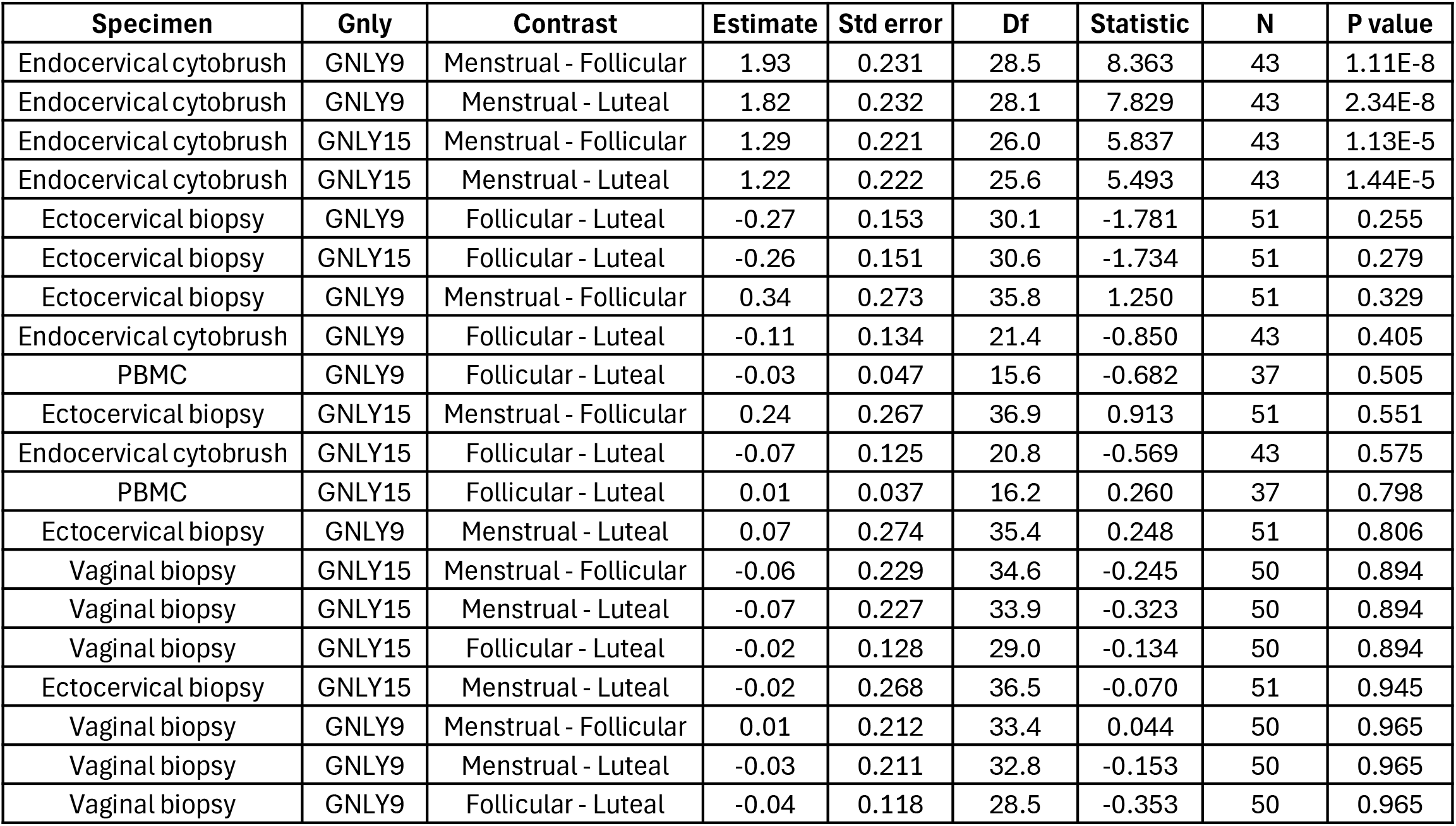
Results of linear mixed effects models comparing the number of GNLY-expressing lymphocytes between phases. . The model fixed effect was menstrual phase, the random effect was participant, and the outcome was the log10 transformed number of GNLY-expressing cells per sample. Estimate indicates the log10 difference in number of GNLY-expressing lymphocytes between the indicated menstrual phases. P-values are adjusted for 3 comparisons (Menstrual vs. follicular, menstrual vs. luteal, and follicular vs. luteal) per specimen and form of GNLY, except for PBMC where no menstrual phase samples were collected and only one comparison was performed.

| Specimen | Gnly | Contrast | Estimate | Std error | Df | Statistic | N | P value |
| --- | --- | --- | --- | --- | --- | --- | --- | --- |
| Endocervical cytobrush | GNLY9 | Menstrual - Follicular | 1.93 | 0.231 | 28.5 | 8.363 | 43 | 1.11E-8 |
| Endocervical cytobrush | GNLY9 | Menstrual - Luteal | 1.82 | 0.232 | 28.1 | 7.829 | 43 | 2.34E-8 |
| Endocervical cytobrush | GNLY15 | Menstrual - Follicular | 1.29 | 0.221 | 26.0 | 5.837 | 43 | 1.13E-5 |
| Endocervical cytobrush | GNLY15 | Menstrual - Luteal | 1.22 | 0.222 | 25.6 | 5.493 | 43 | 1.44E-5 |
| Ectocervical biopsy | GNLY9 | Follicular - Luteal | -0.27 | 0.153 | 30.1 | -1.781 | 51 | 0.255 |
| Ectocervical biopsy | GNLY15 | Follicular - Luteal | -0.26 | 0.151 | 30.6 | -1.734 | 51 | 0.279 |
| Ectocervical biopsy | GNLY9 | Menstrual - Follicular | 0.34 | 0.273 | 35.8 | 1.250 | 51 | 0.329 |
| Endocervical cytobrush | GNLY9 | Follicular - Luteal | -0.11 | 0.134 | 21.4 | -0.850 | 43 | 0.405 |
| PBMC | GNLY9 | Follicular - Luteal | -0.03 | 0.047 | 15.6 | -0.682 | 37 | 0.505 |
| Ectocervical biopsy | GNLY15 | Menstrual - Follicular | 0.24 | 0.267 | 36.9 | 0.913 | 51 | 0.551 |
| Endocervical cytobrush | GNLY15 | Follicular - Luteal | -0.07 | 0.125 | 20.8 | -0.569 | 43 | 0.575 |
| PBMC | GNLY15 | Follicular - Luteal | 0.01 | 0.037 | 16.2 | 0.260 | 37 | 0.798 |
| Ectocervical biopsy | GNLY9 | Menstrual - Luteal | 0.07 | 0.274 | 35.4 | 0.248 | 51 | 0.806 |
| Vaginal biopsy | GNLY15 | Menstrual - Follicular | -0.06 | 0.229 | 34.6 | -0.245 | 50 | 0.894 |
| Vaginal biopsy | GNLY15 | Menstrual - Luteal | -0.07 | 0.227 | 33.9 | -0.323 | 50 | 0.894 |
| Vaginal biopsy | GNLY15 | Follicular - Luteal | -0.02 | 0.128 | 29.0 | -0.134 | 50 | 0.894 |
| Ectocervical biopsy | GNLY15 | Menstrual - Luteal | -0.02 | 0.268 | 36.5 | -0.070 | 51 | 0.945 |
| Vaginal biopsy | GNLY9 | Menstrual - Follicular | 0.01 | 0.212 | 33.4 | 0.044 | 50 | 0.965 |
| Vaginal biopsy | GNLY9 | Menstrual - Luteal | -0.03 | 0.211 | 32.8 | -0.153 | 50 | 0.965 |
| Vaginal biopsy | GNLY9 | Follicular - Luteal | -0.04 | 0.118 | 28.5 | -0.353 | 50 | 0.965 |

**Table S4.** Results of linear mixed effects models comparing the percentage of lymphocytes expressing GNLY between phases. . The model fixed effect was menstrual phase, the random effect was participant, and the outcome was the log10 transformed percent of lymphocytes expressing GNLY. Estimate indicates the log10 difference in percent of GNLY-expressing lymphocytes between the indicated menstrual phases. P-values are adjusted for 3 comparisons (Menstrual vs. follicular, menstrual vs. luteal, and follicular vs. luteal) per specimen and form of GNLY, except for PBMC where no menstrual phase samples were collected and only one comparison was performed.

| **Specimen** | **Gnly** | **Contrast** | **Estimate** | **Std error** | **df** | **Statistic** | **N** | **P value** |
| --- | --- | --- | --- | --- | --- | --- | --- | --- |
| Endocervical cytobrush | GNLY9 | Menstrual - Follicular | 1.13 | 0.303 | 32.5 | 3.73 | 43 | 0.001 |
| Endocervical cytobrush | GNLY9 | Menstrual - Luteal | 1.17 | 0.305 | 31.9 | 3.83 | 43 | 0.001 |
| PBMC | GNLY9 | Follicular - Luteal | 0.07 | 0.039 | 15.6 | 1.68 | 37 | 0.113 |
| PBMC | GNLY15 | Follicular - Luteal | 0.09 | 0.066 | 17.6 | 1.38 | 37 | 0.184 |
| Ectocervical biopsy | GNLY9 | Menstrual - Follicular | 0.19 | 0.117 | 31.6 | 1.66 | 51 | 0.187 |
| Ectocervical biopsy | GNLY9 | Menstrual - Luteal | 0.18 | 0.117 | 31.4 | 1.58 | 51 | 0.187 |
| Endocervical cytobrush | GNLY15 | Menstrual - Luteal | 0.56 | 0.335 | 36.1 | 1.67 | 43 | 0.308 |
| Endocervical cytobrush | GNLY15 | Follicular - Luteal | 0.27 | 0.207 | 23.3 | 1.30 | 43 | 0.308 |
| Endocervical cytobrush | GNLY15 | Menstrual - Follicular | 0.29 | 0.332 | 36.6 | 0.87 | 43 | 0.391 |
| Vaginal biopsy | GNLY15 | Menstrual - Follicular | 0.04 | 0.133 | 35.0 | 0.29 | 50 | 0.770 |
| Vaginal biopsy | GNLY15 | Menstrual - Luteal | 0.07 | 0.132 | 34.2 | 0.54 | 50 | 0.770 |
| Vaginal biopsy | GNLY15 | Follicular - Luteal | 0.03 | 0.075 | 29.1 | 0.44 | 50 | 0.770 |
| Vaginal biopsy | GNLY9 | Menstrual - Follicular | 0.09 | 0.140 | 32.5 | 0.62 | 50 | 0.771 |
| Vaginal biopsy | GNLY9 | Menstrual - Luteal | 0.04 | 0.139 | 32.0 | 0.29 | 50 | 0.771 |
| Vaginal biopsy | GNLY9 | Follicular - Luteal | -0.05 | 0.078 | 28.2 | -0.59 | 50 | 0.771 |
| Endocervical cytobrush | GNLY9 | Follicular - Luteal | 0.04 | 0.182 | 22.3 | 0.21 | 43 | 0.834 |
| Ectocervical biopsy | GNLY15 | Menstrual - Follicular | 0.09 | 0.144 | 33.4 | 0.63 | 51 | 0.844 |
| Ectocervical biopsy | GNLY15 | Menstrual - Luteal | 0.08 | 0.144 | 33.1 | 0.52 | 51 | 0.844 |
| Ectocervical biopsy | GNLY15 | Follicular - Luteal | -0.02 | 0.079 | 29.1 | -0.20 | 51 | 0.844 |
| Ectocervical biopsy | GNLY9 | Follicular - Luteal | -0.01 | 0.064 | 28.5 | -0.14 | 51 | 0.892 |

**Table S5.** Results of linear mixed effects models comparing the GNLY MFI of GNLY-expressing lymphocytes between phases. . The model fixed effect was menstrual phase, the random effect was participant, and the outcome was the log10 transformed GNLY MFI of GNLY-expressing lymphocytes. Estimate indicates the log10 difference in MFIs between the indicated menstrual phases. P-values are adjusted for 3 comparisons (Menstrual vs. follicular, menstrual vs. luteal, and follicular vs. luteal) per specimen and form of GNLY, except for PBMC where no menstrual phase samples were collected and only one comparison was performed.

| Specimen | Gnly | Contrast | Estimate | Std error | df | Statistic | N | P value |
| --- | --- | --- | --- | --- | --- | --- | --- | --- |
| PBMC | GNLY9 | Follicular - Luteal | 0.01 | 0.007 | 15.1 | 2.10 | 37 | 0.053 |
| PBMC | GNLY15 | Follicular - Luteal | 0.02 | 0.016 | 16.5 | 1.38 | 37 | 0.186 |
| Endocervical cytobrush | GNLY9 | Menstrual - Luteal | 0.21 | 0.152 | 25.7 | 1.38 | 32 | 0.296 |
| Endocervical cytobrush | GNLY9 | Follicular - Luteal | 0.14 | 0.108 | 20.7 | 1.33 | 32 | 0.296 |
| Endocervical cytobrush | GNLY15 | Follicular - Luteal | 0.10 | 0.062 | 20.7 | 1.61 | 38 | 0.367 |
| Ectocervical biopsy | GNLY15 | Menstrual - Follicular | 0.06 | 0.065 | 34.1 | 0.99 | 50 | 0.491 |
| Ectocervical biopsy | GNLY15 | Follicular - Luteal | -0.05 | 0.036 | 28.8 | -1.34 | 50 | 0.491 |
| Endocervical cytobrush | GNLY15 | Menstrual - Luteal | 0.09 | 0.097 | 26.6 | 0.97 | 38 | 0.513 |
| Vaginal biopsy | GNLY15 | Menstrual - Follicular | -0.08 | 0.099 | 38.3 | -0.83 | 50 | 0.600 |
| Vaginal biopsy | GNLY15 | Menstrual - Luteal | -0.11 | 0.099 | 37.2 | -1.14 | 50 | 0.600 |
| Vaginal biopsy | GNLY15 | Follicular - Luteal | -0.03 | 0.057 | 30.6 | -0.53 | 50 | 0.600 |
| Vaginal biopsy | GNLY9 | Menstrual - Follicular | -0.04 | 0.055 | 28.1 | -0.77 | 50 | 0.650 |
| Vaginal biopsy | GNLY9 | Menstrual - Luteal | -0.02 | 0.054 | 27.9 | -0.46 | 50 | 0.650 |
| Vaginal biopsy | GNLY9 | Follicular - Luteal | 0.02 | 0.029 | 26.6 | 0.58 | 50 | 0.650 |
| Endocervical cytobrush | GNLY9 | Menstrual - Follicular | 0.07 | 0.150 | 24.1 | 0.44 | 32 | 0.661 |
| Ectocervical biopsy | GNLY9 | Menstrual - Follicular | 0.03 | 0.081 | 40.4 | 0.42 | 48 | 0.676 |
| Ectocervical biopsy | GNLY9 | Menstrual - Luteal | 0.07 | 0.082 | 41.0 | 0.82 | 48 | 0.676 |
| Ectocervical biopsy | GNLY9 | Follicular - Luteal | 0.03 | 0.049 | 31.2 | 0.67 | 48 | 0.676 |
| Ectocervical biopsy | GNLY15 | Menstrual - Luteal | 0.02 | 0.064 | 33.5 | 0.25 | 50 | 0.808 |
| Endocervical cytobrush | GNLY15 | Menstrual - Follicular | -0.01 | 0.097 | 26.9 | -0.06 | 38 | 0.949 |

**Table S6.**
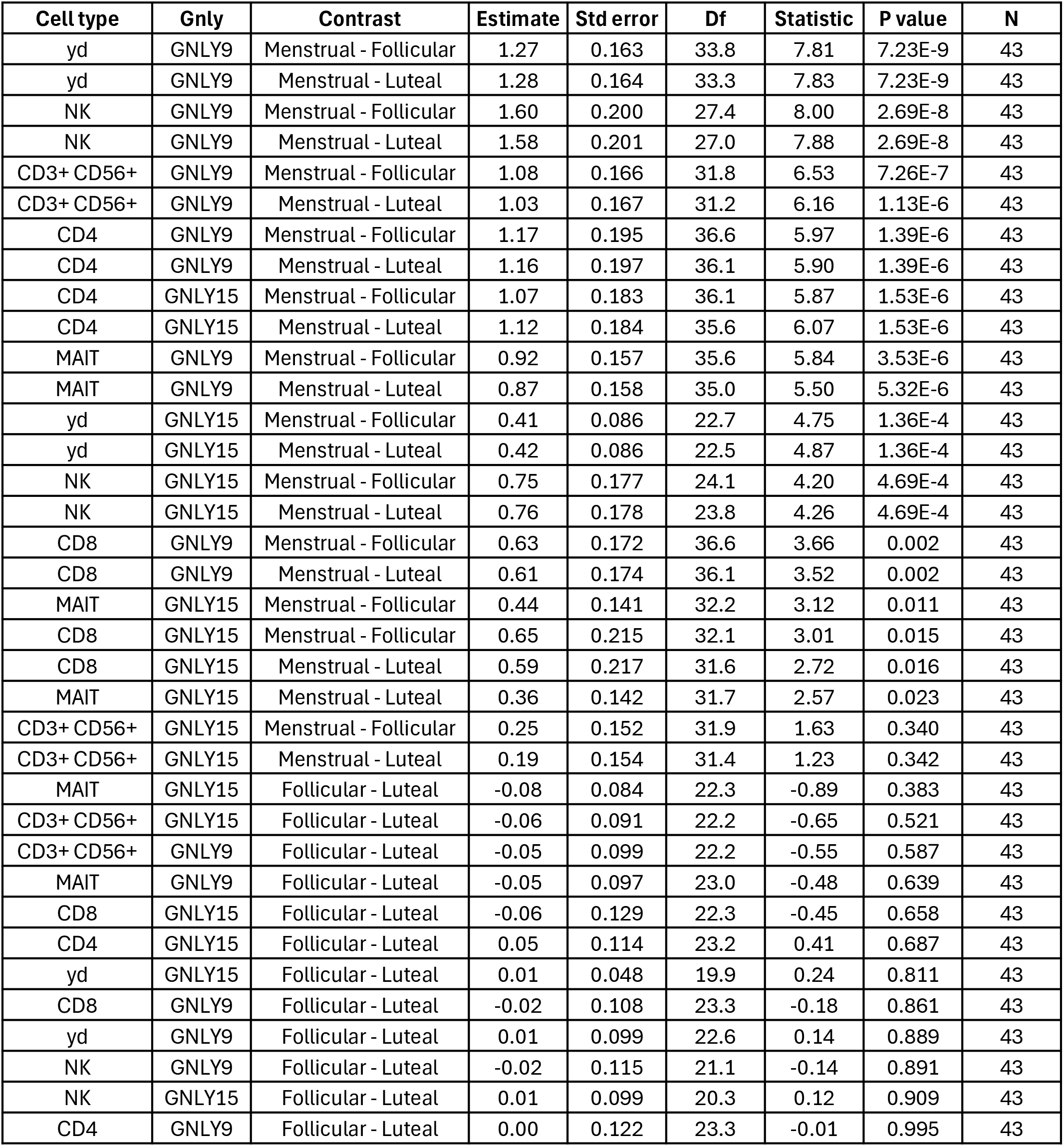
Results of linear mixed effects models comparing the number of GNLY-expressing cells of different cell types between phases in endocervical cytobrushes. . The model fixed effect was menstrual phase, the random effect was participant, and the outcome was the log10 transformed number of GNLY-expressing cells of the indicated cell type per sample. Estimate indicates the log10 difference in number of GNLY-expressing cells between the indicated menstrual phases for the indicated cell type. P-values are adjusted for 3 comparisons (Menstrual vs. follicular, menstrual vs. luteal, and follicular vs. luteal) per specimen, cell type, and form of GNLY.

**Table S7.** Results of linear mixed effects models comparing the percentage of different cell types that express GNLY between phases in endocervical cytobrushes. . The model fixed effect was menstrual phase, the random effect was participant, and the outcome was the log10 transformed percent of GNLY-expression by the indicated cell type. Estimate indicates the log10 difference in percent GNLY-expression between the indicated menstrual phases for the indicated cell type. P-values are adjusted for 3 comparisons (Menstrual vs. follicular, menstrual vs. luteal, and follicular vs. luteal) per specimen, cell type, and form of GNLY.

| Cell type | Gnly | Contrast | Estimate | Std error | Df | Statistic | P value | N |
| --- | --- | --- | --- | --- | --- | --- | --- | --- |
| yd | GNLY9 | Menstrual - Follicular | 1.77 | 0.295 | 18.3 | 6.01 | 1.53E-5 | 35 |
| yd | GNLY9 | Menstrual - Luteal | 1.87 | 0.295 | 18.3 | 6.35 | 1.53E-5 | 35 |
| CD4 | GNLY9 | Menstrual - Follicular | 1.14 | 0.226 | 36.6 | 5.06 | 1.94E-5 | 43 |
| CD4 | GNLY9 | Menstrual - Luteal | 1.15 | 0.228 | 36.1 | 5.05 | 1.94E-5 | 43 |
| CD4 | GNLY15 | Menstrual - Follicular | 1.08 | 0.244 | 36.6 | 4.43 | 1.22E-4 | 43 |
| CD4 | GNLY15 | Menstrual - Luteal | 1.11 | 0.247 | 36.1 | 4.48 | 1.22E-4 | 43 |
| MAIT | GNLY9 | Menstrual - Follicular | 1.38 | 0.329 | 29.3 | 4.20 | 6.76E-4 | 38 |
| MAIT | GNLY9 | Menstrual - Luteal | 1.16 | 0.324 | 27.8 | 3.58 | 0.002 | 38 |
| yd | GNLY15 | Menstrual - Follicular | 0.94 | 0.351 | 18.8 | 2.66 | 0.023 | 35 |
| yd | GNLY15 | Menstrual - Luteal | 0.94 | 0.351 | 18.8 | 2.69 | 0.023 | 35 |
| CD3+ CD56+ | GNLY9 | Menstrual - Follicular | 1.51 | 0.583 | 24.1 | 2.60 | 0.024 | 34 |
| CD3+ CD56+ | GNLY9 | Menstrual - Luteal | 1.53 | 0.571 | 25.1 | 2.69 | 0.024 | 34 |
| NK | GNLY9 | Menstrual - Luteal | 1.41 | 0.559 | 28.5 | 2.52 | 0.053 | 36 |
| CD8 | GNLY9 | Menstrual - Follicular | 0.86 | 0.369 | 33.3 | 2.32 | 0.079 | 40 |
| CD8 | GNLY9 | Menstrual - Luteal | 0.72 | 0.371 | 33.0 | 1.94 | 0.091 | 40 |
| NK | GNLY9 | Menstrual - Follicular | 0.92 | 0.558 | 30.5 | 1.66 | 0.162 | 36 |
| NK | GNLY9 | Follicular - Luteal | 0.49 | 0.376 | 20.6 | 1.29 | 0.210 | 36 |
| MAIT | GNLY9 | Follicular - Luteal | -0.22 | 0.209 | 20.5 | -1.07 | 0.296 | 38 |
| MAIT | GNLY15 | Menstrual - Follicular | 0.63 | 0.478 | 28.3 | 1.32 | 0.423 | 38 |
| MAIT | GNLY15 | Menstrual - Luteal | 0.38 | 0.469 | 26.9 | 0.81 | 0.423 | 38 |
| MAIT | GNLY15 | Follicular - Luteal | -0.25 | 0.301 | 20.2 | -0.83 | 0.423 | 38 |
| NK | GNLY15 | Menstrual - Follicular | 0.44 | 0.566 | 23.6 | 0.77 | 0.501 | 36 |
| NK | GNLY15 | Menstrual - Luteal | 0.68 | 0.556 | 22.0 | 1.23 | 0.501 | 36 |
| NK | GNLY15 | Follicular - Luteal | 0.25 | 0.359 | 18.1 | 0.69 | 0.501 | 36 |
| CD8 | GNLY9 | Follicular - Luteal | -0.14 | 0.237 | 22.2 | -0.58 | 0.569 | 40 |
| yd | GNLY9 | Follicular - Luteal | 0.10 | 0.190 | 16.4 | 0.52 | 0.613 | 35 |
| CD3+ CD56+ | GNLY15 | Menstrual - Follicular | 0.31 | 0.612 | 26.8 | 0.50 | 0.819 | 34 |
| CD3+ CD56+ | GNLY15 | Menstrual - Luteal | 0.40 | 0.597 | 27.7 | 0.67 | 0.819 | 34 |
| CD3+ CD56+ | GNLY15 | Follicular - Luteal | 0.10 | 0.419 | 20.8 | 0.23 | 0.819 | 34 |
| CD4 | GNLY15 | Follicular - Luteal | 0.02 | 0.153 | 23.3 | 0.15 | 0.884 | 43 |
| CD4 | GNLY9 | Follicular - Luteal | 0.01 | 0.141 | 23.3 | 0.06 | 0.949 | 43 |
| CD3+ CD56+ | GNLY9 | Follicular - Luteal | 0.02 | 0.395 | 19.6 | 0.05 | 0.958 | 34 |
| yd | GNLY15 | Follicular - Luteal | 0.01 | 0.227 | 16.6 | 0.04 | 0.972 | 35 |
| CD8 | GNLY15 | Menstrual - Follicular | 0.21 | 0.510 | 33.0 | 0.41 | 0.989 | 40 |
| CD8 | GNLY15 | Menstrual - Luteal | 0.21 | 0.513 | 32.7 | 0.42 | 0.989 | 40 |
| CD8 | GNLY15 | Follicular - Luteal | 0.00 | 0.327 | 22.1 | 0.01 | 0.989 | 40 |

**Table S8.** Univariate linear models assessing associations between participant characteristics and number of luteal phase vaginal swabs containing granulysin. One model was run per variable (left column). The response variable was number of swabs in the luteal phase with detectable granulysin. The response variable was log10 transformed after having 1 added to it (to account for zeros prior to log transformation).

| Variable | Term | Estimate | Standard error | t-value | P-value |
| --- | --- | --- | --- | --- | --- |
| Age | Age | 0.01 | 0.008 | 1.61 | 0.12 |
| HSV-2 | Positive | 0.29 | 0.16 | 1.88 | 0.07 |
| BV | Don't know or prefer not to answer | 0.24 | 0.21 | 1.15 | 0.26 |
|  | Positive | 0.10 | 0.16 | 0.64 | 0.53 |
| Contraception | Yes | -0.13 | 0.14 | -0.88 | 0.39 |
| Lubricant | Yes | -0.27 | 0.15 | -1.82 | 0.08 |
| History of pregnancy and birth | Past pregnancy, gave birth | 0.57 | 0.15 | 3.92 | 0.0009 |
|  | Past pregnancy, no birth | 0.17 | 0.16 | 1.06 | 0.32 |
|  | Missing | 0.26 | 0.20 | 1.31 | 0.20 |

**Table S9.**
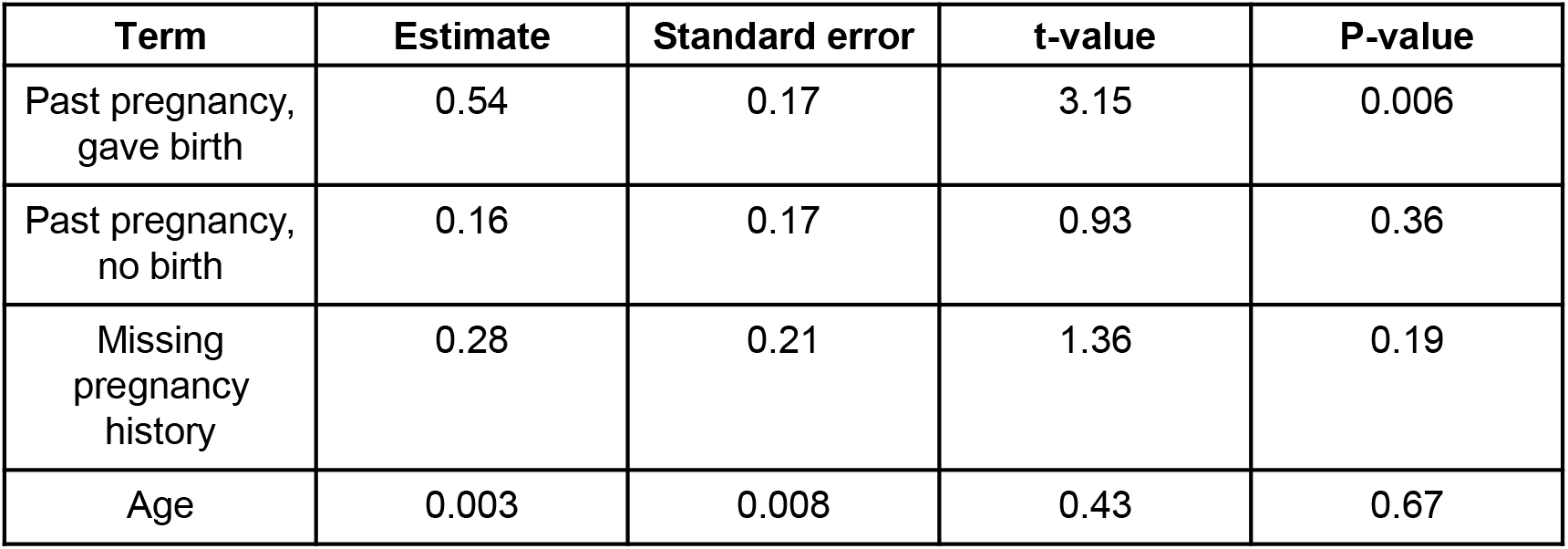
Result of linear model assessing associations between age and history of pregnancy and birth with the number of luteal phase vaginal swabs containing granulysin. The response variable was number of swabs in the luteal phase with detectable granulysin. The response variable was log10 transformed after having 1 added to it (to account for zeros prior to log transformation).

| Term | Estimate | Standard error | t-value | P-value |
| --- | --- | --- | --- | --- |
| Past pregnancy,<br>gave birth | 0.54 | 0.17 | 3.15 | 0.006 |
| Past pregnancy,<br>no birth | 0.16 | 0.17 | 0.93 | 0.36 |
| Missing<br>pregnancy<br>history | 0.28 | 0.21 | 1.36 | 0.19 |
| Age | 0.003 | 0.008 | 0.43 | 0.67 |

